# Establishment and comparative genomics of a high-quality collection of mosquito-associated bacterial isolates - MosAIC (Mosquito-Associated Isolate Collection)

**DOI:** 10.1101/2023.10.04.560816

**Authors:** Aidan Foo, Laura E. Brettell, Holly L. Nichols, 2022 UW-Madison Capstone in Microbiology Students, Miguel Medina Muñoz, Jessica A. Lysne, Vishaal Dhokiya, Ananya Ferdous Hoque, Doug E. Brackney, Eric P. Caragata, Michael Hutchinson, Marcelo Jacobs-Lorena, David J. Lampe, Edwige Martin, Claire Valiente Moro, Michael Povelones, Sarah M. Short, Blaire Steven, Jiannong Xu, Timothy D. Paustian, Michelle R. Rondon, Grant L. Hughes, Kerri L. Coon, Eva Heinz

## Abstract

Mosquitoes transmit medically important human pathogens, including viruses like dengue virus and parasites such as *Plasmodium* spp., the causative agent of malaria. Mosquito microbiomes are critically important for the ability of mosquitoes to transmit disease-causing agents. However, while large collections of bacterial isolates and genomic data exist for vertebrate microbiomes, the vast majority of work in mosquitoes to date is based on 16S rRNA gene amplicon data that provides limited taxonomic resolution and no functional information. To address this gap and facilitate future studies using experimental microbiome manipulations, we generated a bacterial <u>Mos</u>quito-<u>A</u>ssociated Isolate <u>C</u>ollection (MosAIC) consisting of 392 bacterial isolates with extensive metadata and high-quality draft genome assemblies that are publicly available for use by the scientific community. MosAIC encompasses 142 species spanning 29 bacterial families, with members of the *Enterobacteriaceae* comprising 40% of the collection. Phylogenomic analysis of three genera, *Enterobacter, Serratia*, and *Elizabethkingia*, reveal lineages of mosquito-associated bacteria isolated from different mosquito species in multiple laboratories. Investigation into species’ pangenomes further reveals clusters of genes specific to these lineages, which are of interest for future work to identify functions underlying mosquito host association. Altogether, we describe the generation of a physical collection of mosquito-associated bacterial isolates, their genomic data, and analyses of selected groups in context of genome data from closely related isolates, providing a unique, highly valuable resource to investigate factors for bacterial colonisation and adaptation within mosquito hosts. Future efforts will expand the collection to include broader geographic and host species representation, especially from individuals collected from field populations, as well as other mosquito-associated microbes, including fungi, archaea, and protozoa.

## Introduction

Mosquitoes (Diptera: Culicidae) are the major vectors of some of the world’s most important pathogens. The mosquito microbiome can influence all aspects of host biology (1), including the mosquito’s ability to transmit human pathogens (2–9). Given these functions and the increasing resistance of mosquitoes against insecticides (10,11), microbiome manipulation is a promising and increasingly relevant alternative avenue for future applications in vector control. It is thus of urgent relevance to gain a better understanding of the mosquito microbiome’s functional composition and its interactions with the host and invading pathogens.

The composition of the mosquito microbiome is dynamic and affected by host species (12,13) and geography (14,15), and varies both across individuals (16) as well as across an individual’s life stage (17–19). Nevertheless, 16S rRNA gene amplicon sequencing studies have identified a number of commonly present bacterial genera, including *Enterobacter*, *Serratia*, *Asaia*, *Pantoea*, *Elizabethkingia* and *Cedecea* (16,19–21). Results from amplicon sequencing are however limited in taxonomic resolution and provide no functional information (22), rendering it challenging to draw conclusions beyond presence-absence. Experimental manipulations have demonstrated the capacity of specific bacterial isolates to influence mosquito life history in diverse ways. For example, *Serratia marcescens* has been shown to suppress adult feeding behaviour (23,24), while *Asaia* has been shown to accelerate larval development (25,26) and activate mosquito immune genes (27). *Serratia marcescens* and *Enterobacter cloacae* have further been shown to positively influence both adult longevity and egg hatch rates in *Aedes aegypti* (28). However, another study showed *S. marcescens* to have larvicidal properties (29), highlighting the need for strain-level resolution to fully understand mosquito-microbiome interactions.

Understanding why certain bacteria are successful colonisers and how the host and microbial community interact will facilitate development of microbe-based approaches for vector control. This includes strategies like paratransgenesis, where genetically modified bacteria are introduced into the host to express effector molecules to block pathogen transmission or interfere with mosquito longevity or reproduction (30–33). In addition to their medical relevance, mosquitoes are also attractive experimental organisms, as they are amenable to microbiome manipulation through the use of germ-free and gnotobiotic techniques. One such approach involves hatching surface sterilised eggs to produce axenic (microbe-free) larvae that can then be seeded with a microbial community through inoculation of larval water (1,34–36). Further, complete mosquito microbiomes can be successfully extracted, cryopreserved (37), and transplanted between different mosquito species (38). Using these approaches, we recently determined the ability of *Cedecea neteri* to form biofilms, showing this to be a key factor that contributes to colonisation of *Aedes aegypti* mosquitoes (39). We also found the transcriptome of transplant recipients responded similarly when receiving a microbiome from a laboratory-reared donor regardless of mosquito species, but showed more differentially expressed genes when the donor was field-caught (40).

The taxonomic composition of the mosquito-microbiome and its effects on mosquito life history are well characterised in the literature (41–50). However, our understanding of factors that drive bacterial colonisation in the mosquito remain poorly understood. Virulence factors are genes that contribute to bacterial establishment and persistence within a host, including insects (51), and their identification is facilitated by use of whole genome analyses. For example, comparative genomics of *Serratia* isolated from different mosquito species shows differing virulence factor profiles, suggesting differences in their ability to infect the mosquito host (52). Indeed, within other fields, insights into symbiont colonisation and host interactions are facilitated by large-scale dedicated genomic collections of bacterial isolates (53–57)–a resource currently absent in the mosquito microbiome field.

With the aim of facilitating future mosquito microbiome studies by the scientific community at large, we report the generation of a collection of mosquito-associated bacterial isolates with accompanying sample metadata and high-quality assembled genomes. We then use this collection to show the presence of potentially mosquito-associated lineages within three focal genera: *Enterobacter*, *Serratia*, and *Elizabethkingia*. Our results identify highly conserved lineage-specific core genes that represent promising candidates for future studies to characterize functions underlying bacterial interactions with mosquito hosts.

## Results

### 392 draft genomes assembled from the mosquito microbiome and associated environments

MosAIC represents a collection of bacterial isolates and high-quality draft genomic assemblies, including 295 from mosquitoes, 83 from mosquito larval habitats, and 14 from non-mosquito Diptera and their associated environments sampled from both the laboratory and field (Fig. 1). *Aedes aegypti* is the most commonly represented mosquito host species, for which 112 isolates were cultured from eggs, larvae, or adults, followed by *Aedes albopictus* (n=71), *Culex pipiens* (n=23), *Anopheles quadrimaculatus* (n=19), *Aedes triseriatus* (n=12), *Anopheles gambiae* (n=15), and *Toxorhynchites amboinensis* (n=10) (Fig. 1). As it is only female mosquitoes that transmit pathogens, MosAIC is heavily skewed towards female-derived isolates. Of the 245 total isolates recovered from adult mosquitoes, 219 were derived from females. Across all adults, the majority were obtained from non-blood-fed (n=168) as compared to blood-fed females (n=51) (Fig. 1). Adult-derived isolates were also predominantly recovered from whole-body (n=211) as compared to midgut samples (n=90) (Fig. 1).

**Fig. 1:**
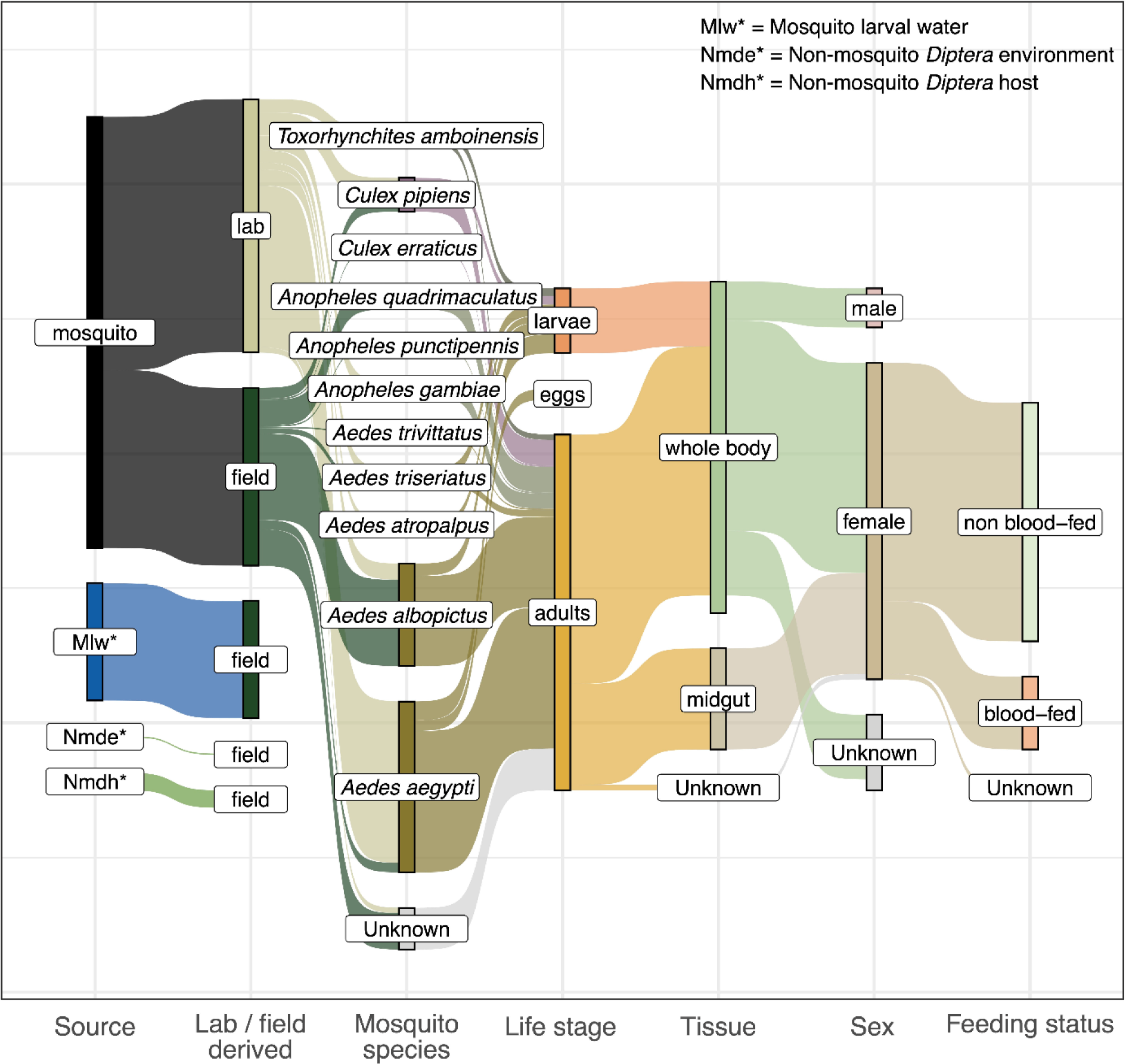
Origin of bacterial isolates in MosAIC. Metadata category names and definitions follow those presented in Table S1. “Unknown” denotes isolates for which a given metadata category is valid but missing. For example, a subset of mosquito samples could not be assigned a species but are derived from adult-stage mosquitoes. Where a given metadata category is invalid, the connection between bars is dropped. For example, feeding status cannot be determined through egg samples.

We sequenced in total 82 GB of read data containing 4.2 megabase pairs (Mbp) on average per sample, ranging between 0.26 Mbp and 27.1 Mbp (Fig. S1). After quality-checking reads, assemblies were assessed based on single-copy core gene content and average genome coverage. Our first filtering threshold (CheckM completeness >98%, CheckM contamination <5%) reduced the dataset to 396 genomes, and after filtering for coverage (>10X), this was finalised to MosAIC’s 392 high-quality assemblies (Figs. 2A-C). The mean completeness score in the collection is 99.67% (98.01 - 100%) and the mean contamination score is 0.59% (0 - 4.40%). Genomes in the collection range in size between 2.21 and 8.38 Mbp, and N50 values range between 9.14 kilobase pairs (KBP) and 3.09 Mbp (Figs. 2D-E). Genome sizes and the number of predicted genes of isolates in the collection are linearly correlated (Fig. S2) (58).

**Fig. 2:**
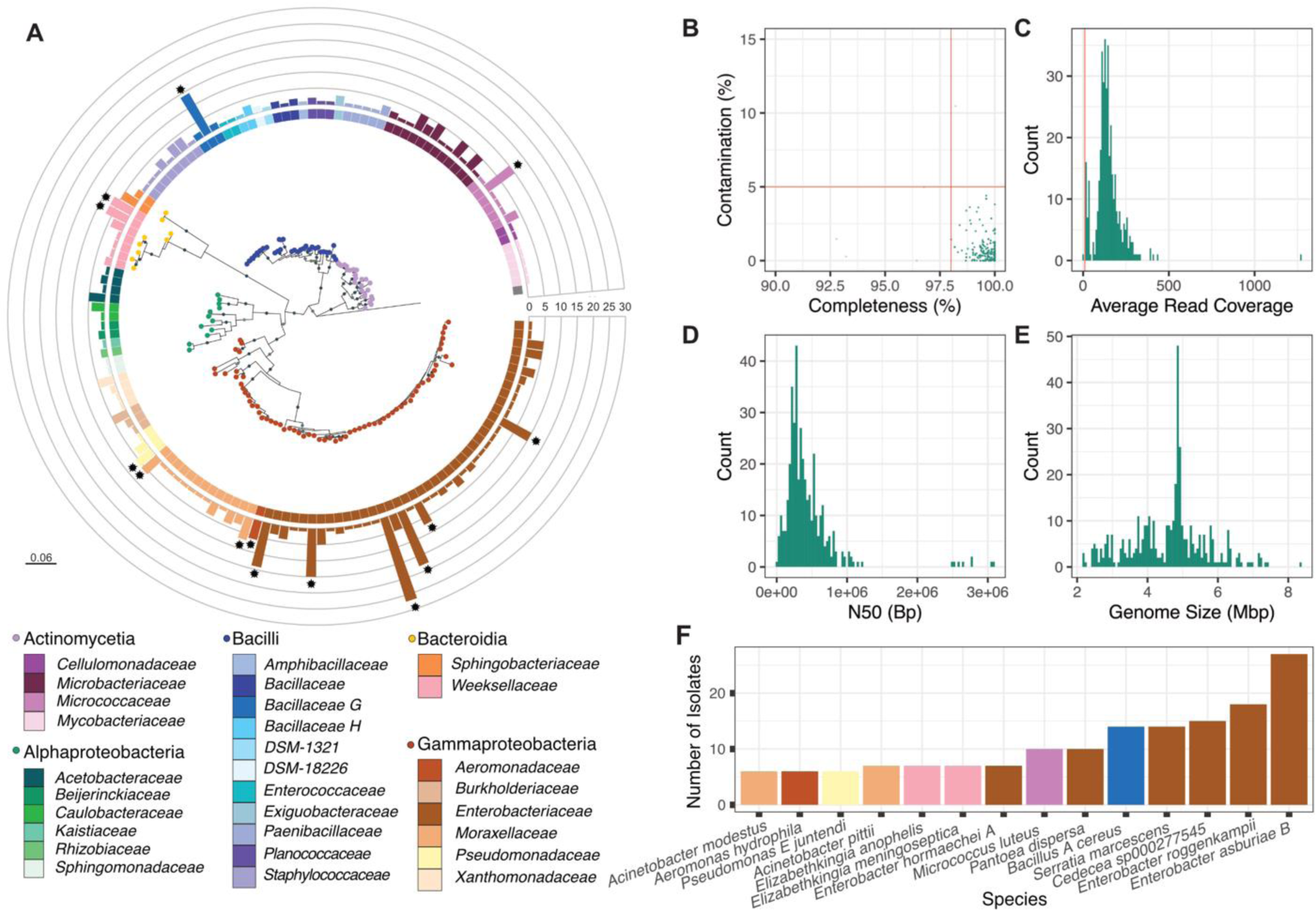
Phylogeny of single species representatives from MosAIC, along with quality-assurance metrics for related genome assemblies. (A) Maximum likelihood tree built using IQ-TREE2 and 16S rRNA gene sequences predicted with Baarnap. Each node is a species representative coloured according to class. Bars at each tip represent the number of isolates present in the species cluster, defined using a secondary clustering threshold of 95% Average Nucleotide Identity (ANI) with dRep. Bars are colour coded according to family information obtained using the Genome Taxonomy Database and classifier GTDB-Tk. Points at the tip of bars delineate highly representative species clusters. Evolutionary scale is displayed on the bottom left of the figure panel. (B) Genome completeness and contamination metrics obtained using CheckM. Each point represents a draft genome assembly. Red lines indicate cutoffs for 98% completeness and 5% contamination. (C) Histogram showing average read coverage reported using QUAST. The vertical red line represents a 10X filter cutoff. (D, E) Histograms showing N50 and genome size across the collection. Green bars represent high-quality genomes within the collection (CheckM completeness >98%, contamination <5%, and >10X coverage). Bp = Base-pairs, Mbp = Mega base-pairs. (F) Highly represented species (>5 isolates) within the collection. Each bar is coloured according to family.

### MosAIC highlights species representatives of known mosquito-associated bacteria

The Genome Taxonomy Database (GTDB) was used to predict the taxonomy of isolates (Table S2). Sixty-two genomes shared >99% ANI with GTDB reference genomes, while 305 shared between 95 - 99% ANI and 25 shared <95% ANI, potentially representing novel species (59). As such, 367 isolates were assigned to a species, with the remaining 25 genomes assigned to their respective genera. Of these 25 genomes, six isolates were assigned to *Leucobacter*, six isolates were assigned to *Microbacterium*, and the remaining 12 isolates were each assigned to individual genera (Fig. S3).

Summarising the collection into species clusters using dRep resulted in 142 single species representatives (Fig. 2A) in 5 bacterial classes: Actinomycetes (4 families), Alphaproteobacteria (6 families), Bacilli (11 families), Bacteroidia (2 families), and Gammaproteobacteria (6 families) (Fig. 2A). The collection is dominated by Gammaproteobacteria; in particular, the *Enterobacteriaceae* comprised 42 species representatives and 157 isolates. The *Microbacteriaceae* and *Weeksellaceae* are also well represented, with 31 and 21 isolates respectively. As such, 6/10 most well-represented species in the collection are *Enterobacteriaceae*, with the remainder belonging to *Bacillaceae*, *Micrococcaceae*, *Moraxellaceae*, and *Weeksellaceae* (Fig. 2F).

### MosAIC isolates are distributed across a range of different sample types

MosAIC is accompanied by comprehensive metadata associated with each isolate (Table S1). While our collection is derived from opportunistic samples and not under a sampling framework, we note that *Acetobacteraceae, Aeromonadaceae* and *Amphibacillaceae* were only isolated from non-blood-fed female and male adult mosquitoes, while *Sphingobacteriaceae* were exclusively isolated from blood-fed females (Fig. S4A). Taxa were well represented from both laboratory- and field-derived mosquitoes from adult, larvae, larval water, and egg samples, but there were differences between the overall number of isolates between these two sources. For example, 78% (n=21) of isolates assigned to the bacterial family *Bacillaceae* were from field-derived samples, whereas 84% (n=26) of isolates assigned to the *Microbacteriaceae* were from laboratory-derived samples (Fig. S4B). In contrast, the most dominant family represented in the collection, *Enterobacteriaceae,* was split relatively equally with 48% (n=75) of isolates assigned to this family originating from field-derived samples and 52% (n=82) from laboratory-derived samples.

We also note interesting patterns amongst genera within the *Enterobacteriaceae* – the most abundant and frequently observed family across MosAIC. Of laboratory- and field-derived isolates, we note 91% (n=31) of *Pantoea* isolates were from field-derived samples, whereas 80% of *Enterobacter* (n=44) isolates were laboratory-derived (Fig. S5A). *Atlantibacter*, *Cedecea*, *Chania*, and *Rouxiella* were exclusively laboratory-derived (Fig. S5A). 57% (n=90) of *Enterobacteriaceae* were isolated from female adult mosquitoes, while only *Pantoea* and *Enterobacter* were isolated from both male and female adult mosquitoes (Fig. S5B).

### Mosquito-associated bacteria contain known and predicted virulence factors

We screened the collection for known and predicted virulence factor genes using the virulence factor database (VFDB). Gene hits were categorised using the VFDB scheme and therefore divided into the VFDB categories adherence, antimicrobial activity/competitive advantage, biofilm, effector delivery system, exoenzyme, exotoxin, immune modulation, invasion, motility, nutritional/metabolic factor, regulation, stress survival and other (Fig. 3).

**Fig. 3:**
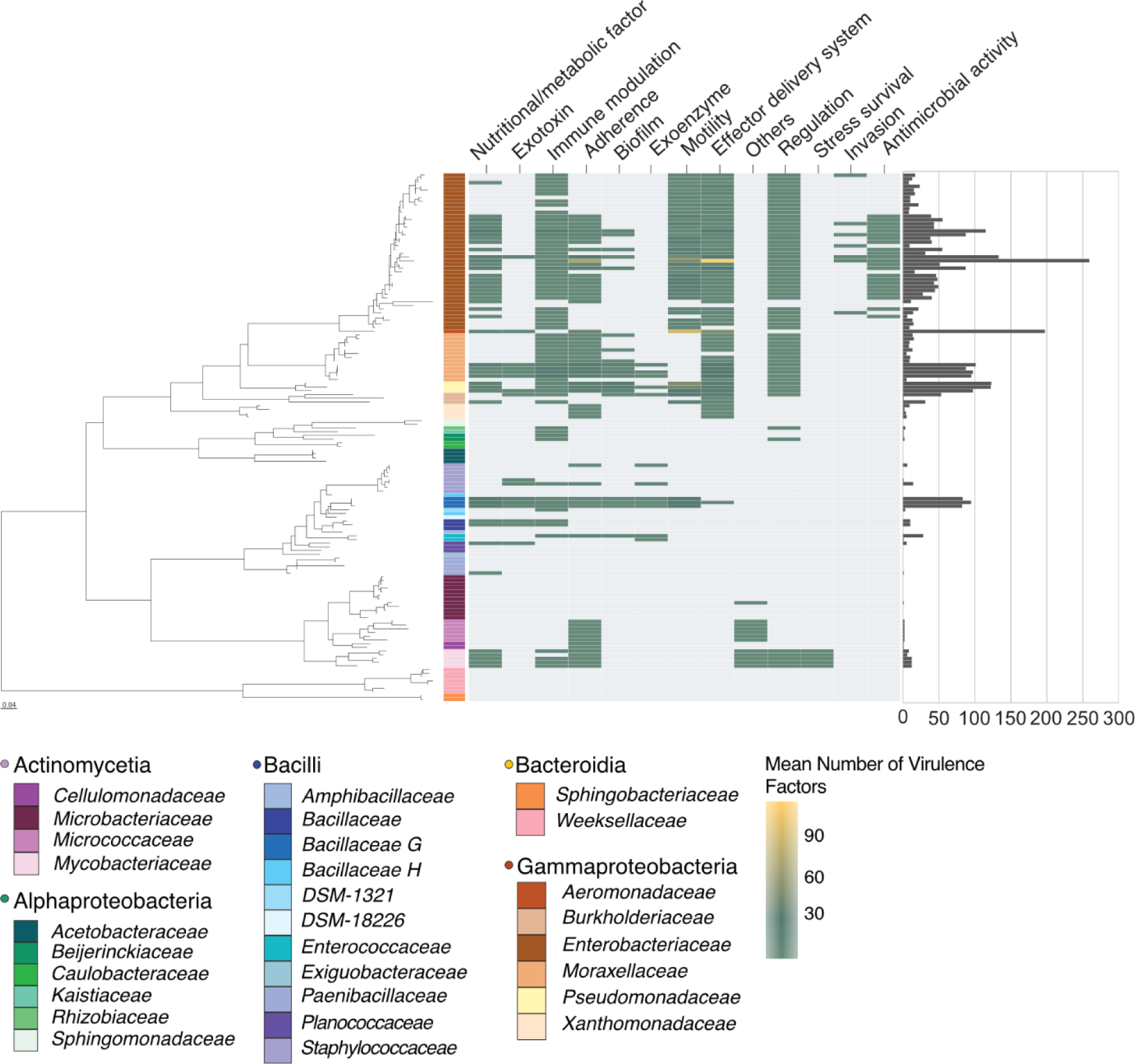
Heatmap of the distribution of virulence factors across all MosAIC genomes. Genes fall within one of 13 different categories (*top*). The guidance tree on the left is a maximum likelihood tree built using IQ-TREE2 and Baarnap-predicted 16S rRNA gene sequences from species clusters defined with dRep. Tiles denote the mean number of virulence factor genes identified within a given species cluster, following a gradient from blue (low) to yellow (high). Grey tiles denote species clusters for which zero predicted virulence factor genes were identified. Bacterial families are colour-coded in the figure legend. The bar chart on the right shows the total number of genes identified within each species cluster.

In total, we identified 11,774 virulence factor genes across the collection and 1,203 uniquely present genes (Table S3). The mean number of virulence genes was highly variable among bacterial classes ranging from Gammaproteobacteria and Bacilli, which contained on average 45 and 52 virulence genes, respectively, to Actinomycetes and Alphaproteobacteria, which contained on average three and one virulence genes, respectively, and Bacteroidia, in which we did not detect any virulence genes (Fig. 3). The distribution of virulence factor categories also varied. For example, stress survival genes were found only in the Actinomycetes. Similarly, genes involved in invasion and antimicrobial activity were only detected in the Gammaproteobacteria. Within bacterial families there was also variation in virulence factor profiles; genes involved in stress survival were only detected in the *Mycobacteriaceae*, and genes involved in invasion and antimicrobial activity were common in the *Enterobacteriaceae* but were not detected in any other family (Fig. 3). On the other hand, genes implicated in roles such as adherence and nutritional/metabolic factors were widespread and present in multiple families. It must be noted, however, that detection of these virulence factors is constrained by the inherent bias of the database composition in that there is limited representation of certain bacterial classes, including a complete absence of genes in the Bacteroidia (Fig. S6).

### Placement of mosquito-associated isolates into wider population structures

To determine the placement of MosAIC isolates in their wider population structures, we selected three taxonomic groups for further analysis (*Enterobacter, Serratia*, and *Elizabethkingia anophelis*), owing to their high representation within the collection, biological interest amongst the mosquito-microbiome field (34), and good understanding of their genetic diversity and environmental / host-associated niches within a population (*i.e.*, their population structure) (60–62). In total, 55 *Enterobacter* isolates, 16 *Serratia* isolates, and 7 *El. anophelis* isolates were phylogenetically placed into their respective populations (Fig. S7-9), which comprise previously sequenced isolates from a variety of environmental niches. The placement of these mosquito-associated genomes within their larger species clades helped refine GTDB-taxonomic classifications; *Serratia nevei* and *Serratia bockelmanni* are now within the *S. marcescens* clade (Fig. 4B), and a single *Enterobacter hormaechei* isolate falls within the *Enterobacter xiagfangensis* clade (Fig. 4A).

**Fig. 4:**
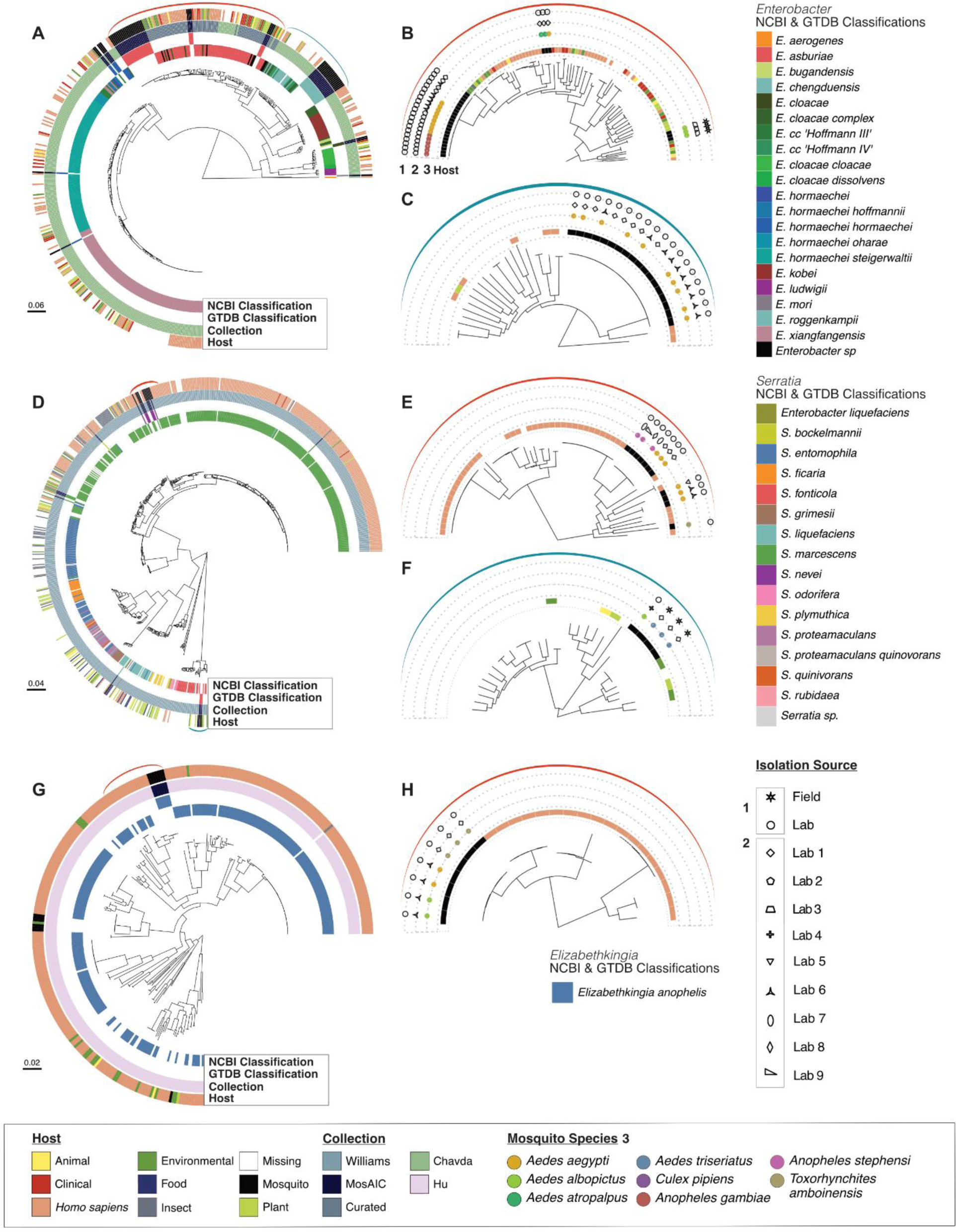
Selected genus population structures with improved mosquito representation. Population structures based on genomic collections for (A-C) *Enterobacter* (61), (D-F) *Serratia* (62), and (G, H) *Elizabethkingia anophelis* (60), with added mosquito-derived representation from MosAIC and a manually curated set of *En. asburiae* genomes. Phylogenies were built using a maximum likelihood approach within IQ-TREE2 (63) and 1000 bootstraps, using SNP-filtered core gene alignments generated with Panaroo (64) and SNP-sites (65). To the right of each population tree are subsets highlighting mosquito-associated lineages within a population, with the coloured brackets highlighting their location within a given population tree. The rings of each population phylogeny (A, D, G) denote, from inner to outer, the NCBI classification from the original study, GTDB classifications from MosAIC, genomic collection, and host. The subset phylogenies show four rings, which denote: host = the isolate source, 1 = lab or field-derived mosquito, 2 = the laboratory group the isolate was sourced from, and 3 = mosquito species the isolate was cultured from. *Enterobacter liquefaciens* within the *Serratia* phylogeny are derived from (62) and have since been reclassified as *Serratia liquefaciens*.

Population structures revealed the clustering of mosquito-associated isolates within the *Enterobacter* and *Serratia* genera (Figs. 4A, D). For *Enterobacter*, the majority of mosquito-associated isolates grouped with *Enterobacter asburiae* and *Enterobacter rogennkampii,* with MosAIC isolates forming distinct lineages of 21 and 18 isolates, respectively (Figs. 4B-C). Similarly, within *Serratia*, most mosquito-associated isolates belonged to *S. marcescens and S. fonticola*, with seven and three isolates, respectively (Figs. 4E-F). Within species, mosquito-associated isolates are often clustered together. The largest mosquito-associated lineage of *En. asburiae* contained isolates from five different mosquito species (*Ae. aegypti*, *Ae. albopictus, Aedes atropalpus, An. gambiae, Cx. pipiens*) and two different, geographically distinct, laboratories (Fig. 4B). Similarly, the *En. roggenkampii* mosquito-associated lineage consisted of isolates from laboratory-reared mosquitoes from two laboratories (Fig. 4C). In the major *S. marcescens* mosquito-associated lineage, there are seven isolates derived from two mosquito species (*An. stephensi, Ae. aegypti*) and five different laboratories (Fig. 4E). Of these seven isolates, five are from MosAIC and two originate from a previous study (52). Interestingly, other *Serratia* isolates including *S. bockelmannii* are derived from water and insect-associated sources and are distantly related to mosquito-associated lineages (Fig. 4D). A similar observation can be seen for *El. anophelis*, where MosAIC-derived isolates from three different mosquito species and two different laboratories clustered together (Figs. 4G-H).

Not all mosquito-associated isolates formed distinct within-species clades. Particular *S. marcescens* (Fig. 4E) and *S. fonticola* clades (Fig. 4F) form multiple lineages reflecting their laboratory sources. However, these splits include singleton lineages and therefore may represent species where the full diversity has yet to be sampled. *El. anophelis* from MosAIC form a distinct mosquito-associated lineage and do not cluster with the previously sequenced mosquito-derived *El. anophelis* (Figs. 4G-H).

### Genetic diversity of pangenome gene classifications of mosquito-associated bacteria

To investigate the conservation of genes within mosquito-associated bacteria, we explored the genetic diversity of *En. asburiae*, *S. marcescens*, and *El. anophelis.* The core genome of each species (gene frequency >95% across all within-species lineages) amounted to 3 109 genes in *En. asburiae*, 2 443 genes in *S. marcescens*, and 1 836 genes in *El. anophelis* (Figs. 5A-C). Gene accumulation curves showed the addition of subsequent genomes did not lead to a plateau of genetic diversity, indicating that all pangenomes analysed were open (Fig. S10).

**Fig. 5:**
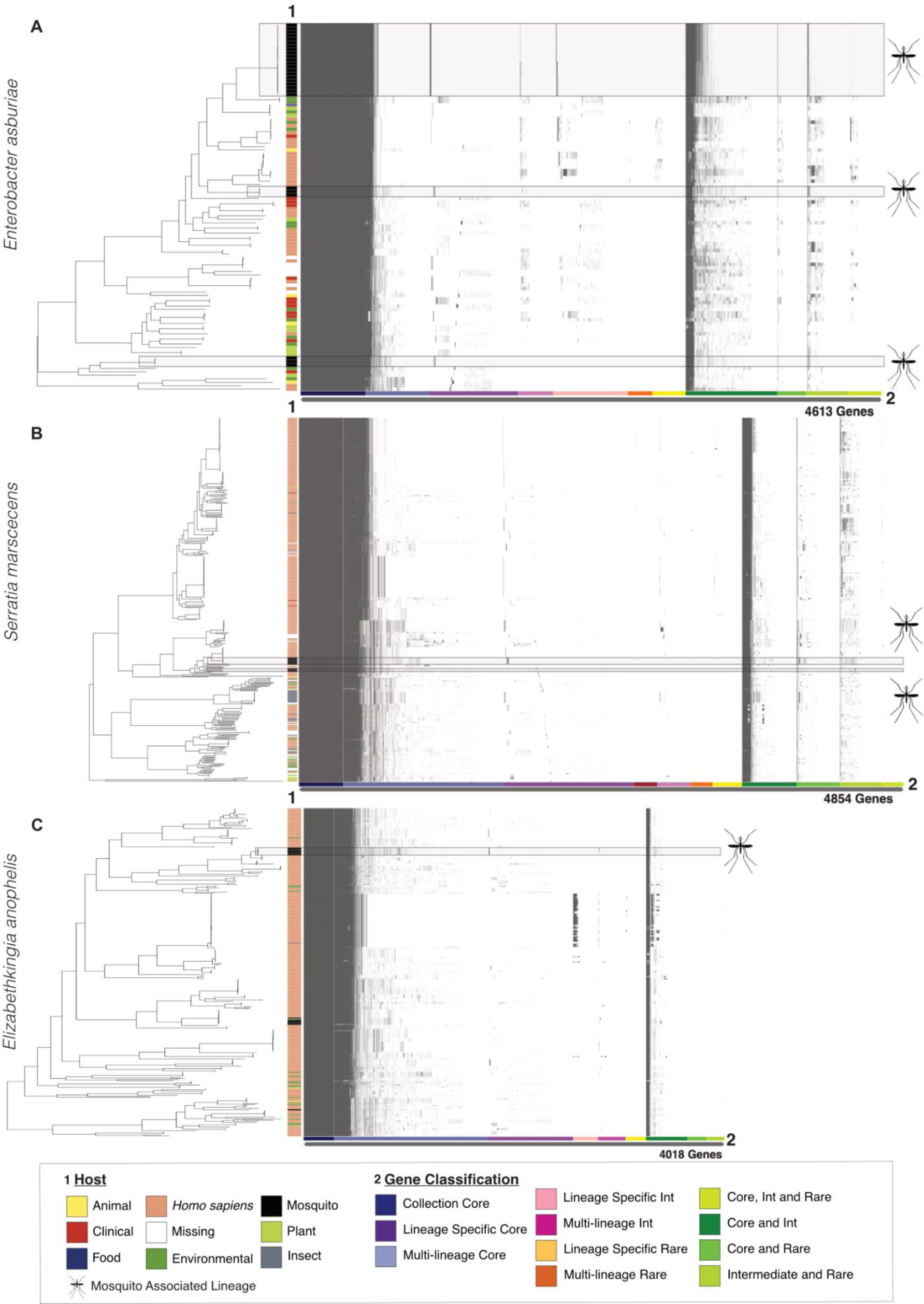
Pangenomes of *Enterobacter asburiae, Serratia marcescens*, and *Elizabethkingia anophelis* with highlighted mosquito-associated lineages. Panels (A-C) depict gene presence / absence within each species, generated with Panaroo (64). Phylogenies and matrices are shaded grey to highlight mosquito-associated lineages defined by PopPUNK (66). The *y*-axis shows the host each bacterium was isolated from, denoted as 1 Host in the figure legend. The *x*-axis shows sub-classifications of the accessory genome, denoted as 2 Gene Classification in the figure legend. Here, subclassification was performed using the twilight package (67). In brief, the classification of each gene was first defined by determining their frequency within a lineage (Core, genes present in ≥95% of strains in a lineage; Int, genes present in ≥15% and ≤95% of strains; Rare, genes present in ≤15% of strains). The resulting gene classifications were then compared across each lineage using genome clusters defined with PopPUNK, which correspond to predicted lineages within the phylogeny (Collection core, genes core to the whole phylogeny; Lineage specific core, genes core to a single lineage; Multi-lineage core, genes core to ≥2 lineages). Genes defined by different classifications across lineages are given a combined class denoted by the green shading.

By analysing the accessory gene content and nucleotide divergence with PopPUNK, we defined six genome clusters that were specific to mosquito-associated isolates–three in *En. asburiae* (Fig. S11), two in *S. marcescens* (Fig. S12), and one in *El. anophelis* (Fig. S13), which may suggest bacterial speciation or niche adaptation to the mosquito host. These lineage-specific core genes were defined as being both specific to a lineage and present at >95% frequency across that single lineage. Within *En. asburiae*, the three mosquito-associated lineages contained 62, 41, and 43 lineage-specific core genes; the two *S. marcescens* mosquito-associated lineages contained 78 and 13 lineage-specific core genes, and the single *El. anophelis* mosquito-associated lineage contained 38 lineage-specific core genes (Figs. 5A-C, Table S4). Interestingly, within *En. asburiae* there were two genes conserved across two different mosquito-associated lineages. These corresponded to a domain of unknown function, DUF4224 (UniRef90 A0A156GGP5), shared between lineages 1 and 8, and an HNH endonuclease (UniRef90 IPI00187EE547) shared between lineages 7 and 8. Lineage-specific intermediate genes (between 15-95% frequency) can be indicative of recently acquired or lost genes within a lineage. *Enterobacter asburiae* contained on average 33 and 2 intermediate core genes within lineages 1 and 8 respectively and the single *El. anophelis* lineage contained 20 intermediate core genes.

The genes within the mosquito-associated clusters have a variety of annotations (Fig. S14). Within *En. asburiae,* the most common annotations across all mosquito-associated isolates were “lipoprotein” (n=84 annotations), “AAA family ATPase” (n=66), “DNA helicase” (n=42), and “phage protein” (n=42). Common annotations for *S. marcescens* were “ATP-binding protein” (n=16 annotations), “response regulator” (n=14), and “restriction endonuclease subunit S (n=14), while common annotations for *El. anophelis* were “translocated intimin receptor (TIR)” (n=21), “phage protein” (n=14), and “A-deaminase” (n=14). In total, we found six annotations shared between *En. asburiae* and *S. marcescens*, five annotations shared between *El. anophelis* and *En. asburiae*, and one annotation shared between *El. anophelis* and *S. marcescens* (Fig. S15). These included specific “TIR” annotations shared between *El. anophelis* and *S. marcescens*, a “HTH luxR-type domain” annotation shared between *El. anophelis* and *En. asburiae*, and a phage-associated “tail fibre assembly protein” annotation shared between *En. asburiae* and *S. marcescens* (Fig. S15).

## Discussion

The mosquito microbiome influences multiple facets of mosquito life history, including the ability of certain species to transmit human pathogens (5). However, while current research in the mosquito microbiome field has greatly improved our understanding of the composition of mosquito-associated bacterial communities and their broad effects on different mosquito phenotypes, our genomic understanding, and therefore the predicted function of specific community members and assemblages, remains poorly understood. Here, we present a collection of 392 mosquito-associated bacterial isolates and their genome sequences, representing the first large-scale community effort to establish a genomic isolate collection for mosquitoes.

Currently, the GTDB (Release 07-RS207) holds 35 mosquito-associated bacterial genomes across 22 species. MosAIC expands this representation to 142 species of mosquito-associated bacteria, acquired from different vector species and life stages and from individuals with different feeding statuses (*i.e.*, blood-fed vs. non-blood-fed adult females) (21,68–70). This collection of isolates can be used to extend previous findings from amplicon-based studies (71–73) using a genotype to phenotype framework. For example, genomic investigation and functional validation of siderophore biosynthetic gene clusters (44,74) or hemolysins (75) in *Serratia* and *Elizabethkingia* may reveal why they are frequently identified post-blood meal (76–79). Furthermore, many bacteria known to be commonly associated with mosquitoes, including *Enterobacter* (28,42)*, Cedecea* (39), *Pantoea* (80,81)*, Elizabethkingia* (76,82–85)*, Aeromonas* (86), and *Acinetobacter* (87) are well represented in the collection. Since the collection of these isolates was not systematic, we did not use statistical inference to indicate differences between isolates and sources but instead pointed to key observations within the collection. An example here are the *Acetobacteraceae* which were abundant and exclusive to non-blood-fed female and male adult mosquitoes compared to blood-fed females, which could reflect their role as sugar oxidisers (88,89). Moreover, as with other host-associated genome collections (54,56,90,91), in compiling MosAIC we discovered 15 genera with no species representation in the GTDB– highlighting the undiscovered taxonomic diversity of mosquito-associated isolates. Now, further work is required to phenotypically characterise these bacteria, including characterising their morphology and growth requirements, determining their genetic content, and elucidating their specific interactions with the mosquito host and other members of the mosquito microbiome.

Mosquitoes are holometabolous insects that undergo complete metamorphosis as they develop and transition from aquatic to terrestrial habitats. During iterative phases of growth and moulting during larval stages and metamorphosis from the larval to adult stage (via a pupal stage), the microbiome is reassembled–resulting in altering composition across life stages (17,77,92,93). Furthermore, the microbiome is highly variable between mosquito populations (1,18,77,94–98). This suggests mosquitoes harbour transient bacteria, similar to butterflies and moths (99,100), rather than specialised bacteria found in social insects such as termites and honey bees that live in colonies comprised of large numbers of related individuals (101,102). In our results, we observed lineages of solely mosquito-associated isolates within populations of three taxa: *Serratia, Enterobacter*, and *El. anophelis*. These mosquito-associated lineages comprised isolates derived from different mosquito species and geographically distinct locations, suggesting adaptation to the mosquito environment. Furthermore, we identified the presence of highly conserved genes within these lineages and in *En. asburiae,* we saw genes which were common to multiple mosquito-associated lineages, suggesting they are beneficial in a mosquito-associated lifestyle. Many of the identified genes implicated in this adaptation remain to be characterised; by annotation using multiple databases, many are described as encoding hypothetical proteins and thus are interesting candidates for further functional investigation. Other genes identified across multiple mosquito-associated lineages were annotated as phage-derived sequences, which may contribute to the introduction of novel accessory elements or expression of chromosomal genes (103). Indeed, this signal may have arisen as a result of laboratory rearing, whereby bacteria have evolved through repeated exposure to the mosquito within a contained insectary environment (104). However, investigating the genetic changes that result from mosquito adaptation is critical for the aim of manipulating these bacteria in the mosquito. To expand on these promising findings, further sampling of mosquito isolates from geographically distinct, field-derived mosquitoes and functional validation of these genes is necessary.

Mosquito-associated bacteria are acquired primarily through horizontal transmission from water to larvae, where they colonise multiple organs, primarily the midgut, with some microbiome members persisting until adulthood (21,79). Within the mosquito, bacteria must withstand a fluctuating environment, including spatial permutations during metamorphosis (93), changes in nutrition (105,106), temperature (107), pH (108), oxidative stress (109), and competition with other members of the microbiota (23,84). The mechanism of adaptation to both an aquatic lifestyle and colonisation in the mosquito remains poorly understood. As such, we screened MosAIC for virulence factors to gain insight into potential colonisation factors and survival strategies amongst these bacteria. This revealed multiple different virulence factor genes, including those involved in adhesion, motility, and biofilm formation, giving a foundation to characterise their function in free-living and host-associated lifestyles. For example, adhesion may be important for horizontal transmission of mosquito symbionts. Within sponges, the vertically transmitted *Ca. Synechococcus spongiarum* lacks type IV pilus genes, while they are present in horizontally transmitted *Ca. Synechococcus feldmannii* (110). The lack of stable transmission over generations likely necessitates alternative attachment mechanisms compared to vertically transmitted symbionts. Adhesion is also likely to play a role in withstanding peristaltic movement during mosquito feeding (111), as seen in other invertebrate systems (112), and specialised motility may be required for movement through the mucosal environment of the mosquito digestive tract, similar to *Burkholderia* symbionts in the *Riptortus pedestris* bean bug (113). It will also be important to further investigate differences between virulence factors in field and laboratory-derived mosquito isolates; for example, flagellar motility is lost in *Acetobacter* from lab-reared *Drosophila* after their close association with the fly through multiple generations, whereas they are retained in field-derived isolates originating from heterogeneous populations (104).

Virulence factor representation overall varied considerably amongst specific families of mosquito-associated bacteria; virulence factor genes were absent within the *Bacteroidia* due to no representation in the VFDB and a heavy skew is observed toward the *Enterobacteriaceae*. This stems from our understanding of virulence factors being heavily influenced by research into human pathogens, many of which fall within the *Enterobacteriaceae* (114), and supports the need for more active research of the roles of these genes within the mosquito (51). Further work to understand the expression, function, and diversity of these genes, as well as their role in free-living and mosquito-associated stages, is paramount for understanding symbiont colonisation and interactions with the mosquito.

The community-based approach of creating MosAIC resulted in a collection of isolates from a diverse range of mosquito-associated environments. However, we acknowledge the biases of this method, including a lack of systematic sampling. The culturing of isolates has resulted in exclusively aerobic bacteria in the collection, and bacteria were isolated on a limited selection of growth media, meaning a potential underrepresentation of strains requiring specific nutrients or growth conditions. This collection also lacks interkingdom representation of fungi, archaea, and protozoa; such will be the target of future sampling efforts. We intend to supplement MosAIC with additional targeted sampling and through the inclusion of metagenomic data (115). This will limit sampling biases through sequencing potential taxa not yet amenable to culturing (116,117) and provide a snapshot of predicted community function to supplement information from single isolates. Given the inherent taxonomic variability of mosquito microbiomes, this gives a compelling reason to address whether community function, rather than taxa, is a conserved unit of the microbiome (118) (*i.e.*, the “song not the singer” hypothesis), as has been postulated in other systems (119–122).

### Conclusion

MosAIC marks a significant milestone as the first large-scale repository of physical isolates and high-quality genomic data associated to the mosquito-microbiome. Using this resource, we have begun to understand the adaptive traits of mosquito-associated symbionts by observing clusters of mosquito-associated bacterial lineages with conserved genes. Notably, we find shared genes between mosquito-associated lineages of *En. asburiae*, suggesting evolutionary convergence driven by mosquito association. To gain further insight of bacterial colonisation within this dynamic system, we emphasise probing the roles of virulence factors in these bacteria; understanding the function and expression of these genes will bring insight to their importance during mosquito development and horizontal transmission from a free-living to mosquito-associated lifestyle. Lastly, we aim to expand this collection through greater systematic sampling from field-derived, disease-endemic sources and parallel metagenomic-based sequencing efforts. This will allow us to build on our population-based observations of mosquito-adaptation to include statistical-based inferences (123) and explore community function. Addressing these questions will bring us a step closer to novel microbe-based mosquito control tools to positively impact public health and vector control efforts.

## Materials and methods

### Sample collection

#### Bacterial isolations and collection curation

Aiming to curate a collection of bacterial isolates that encompasses the diversity of the mosquito microbiome, isolates were recovered from a diversity of mosquito-associated environments (Fig. 1) using a range of media and growth conditions (Table S1). A total of 372 isolates were curated for the collection at the University of Wisconsin-Madison (UW-Madison) – 81 from internal collections comprised of isolates derived from laboratory and field samples, 172 from samples provided by external contributors, and 119 from samples processed by students enrolled in MICROBIO 551 (Capstone Research Project in Microbiology, 2 credits) during the Spring 2022 semester (Table S1). A total of 48 isolates were curated from the collection at the Liverpool School of Tropical Medicine (LSTM), from internal collections comprised of isolates derived from laboratory samples (Table S1).

Isolates derived from existing collections at UW-Madison and LSTM were streaked from frozen glycerol stocks onto agar plates and incubated overnight. Independent bacterial colonies were then inoculated into 3 ml of liquid broth in 14-ml round-bottom tubes (Fisher Scientific, Hampton, NH, USA), incubated overnight with shaking at 200 rpm, and used to generate new glycerol stocks (1 ml culture + 1 ml 40% sterile glycerol) stored in duplicate at −80 °C. Specific incubation temperatures and growth media used follow those provided in Table S1.

Samples from external contributors were received by UW-Madison as either (*i*) agar stabs or liquid cultures of pure isolates inoculated directly from frozen glycerol stocks, or (*ii*) mixed lawns of bacteria on agar plates spread with whole mosquito homogenates. Isolates received as agar stabs or liquid cultures were immediately processed by streak-plating to ensure purity prior to long-term cryopreservation as described above. Samples received as mixed bacterial lawns were scraped into 1 ml of sterile phosphate-buffered saline (PBS, 1X), resuspended, serially diluted and plated on agar prior to incubation for up to 72 hours. Single colonies were then picked for streak-plating prior to long-term cryopreservation of different morphotypes as described above. Specific incubation temperatures and growth media used follow those provided in Table S1.

UW-Madison “Capstone in Microbiology” students isolated bacteria from samples of adult females, larvae, and larval water from internally maintained laboratory cultures of *Ae. aegypti*, *Aedes triseriatus*, *Culex pipiens*, and *Toxorhynchites amboinensis*. Adult female mosquitoes were collected via aspiration, cold anesthetized at −20 °C for 10 minutes, and surface-sterilized with 70% ethanol for 5 minutes followed by three rinses with sterile PBS (1X) or water. Pools of 10 adult females were then combined into a pre-sterilized 2 ml screw cap tube (Sarstedt, Inc., Newton, NC, USA) containing 500 µl PBS and one 5 mm steel bead (Qiagen, Hilden, Germany) prior to bead-beating for 20 seconds to disrupt the cuticle. Cohorts of both sugar-fed and blood-fed adult females were collected for *Ae. aegypti*, while only sugar-fed adult females were collected for *Ae. triseriatus, Cx. pipiens,* and *Tx. amboinensis*. Adult female *Ae. aegypti* were blood fed using defibrinated sheep blood via an artificial membrane feeder. Larvae were collected as fourth instars, surface-sterilized and rinsed prior to homogenization of pools of 10 larvae in 1.7 ml snap-cap tubes (Corning, Inc., Corning, NY, USA) containing 500 µl sterile PBS (1X), using a sterile plastic pestle (Bel-Art Products, Inc., Wayne, NJ, USA). Larval water from trays containing fourth instar larvae was collected in 50 ml screw-cap tubes (Thermo Fisher Scientific, Waltham, MA, USA), centrifuged for 5 minutes at 8,000 x *g*, and resuspended in 2 ml PBS. Resulting homogenates were immediately diluted to extinction in YTG broth (up to 10^-5^ in 96-well plates) and observed for growth for 21 days at 30°C. Wells showing growth were then struck onto YTG agar for isolation as described above.

#### DNA extraction and sequencing

DNA was extracted from each sample using the DNeasy Blood and Tissue kit (Qiagen, Hilden, Germany), following manufacturer’s conditions, with those samples extracted at UW-Madison (Table S1) being subjected to the recommended pre-treatment step for isolation of Gram-positive bacteria, and the samples prepared at LSTM being subjected to the recommended pre-treatment step for Gram-negative bacteria. DNA extracted at LSTM was quantified using a Qubit fluorometer (Thermo Fisher Scientific, Waltham, MA, USA) and used to generate sequencing libraries using the NEBNext Ultra II FS Library Prep kit (New England Biolabs, Ipswich, MA, USA), followed by sequencing on the Illumina MiSeq platform to generate 250 bp paired end reads. DNA extracted at UW-Madison was quantified using a Quantas fluorometer (Promega, Madison, WI, USA) and sent to the UW-Madison Biotechnology Center Next Generation Core facility for library preparation using the Celero EZ DNA-Seq Library Prep kit (Tecan, Männedorf, Switzerland), followed by sequencing on the Illumina NovaSeq6000 platform to generate 150 bp paired end reads.

### Bioinformatics

#### Quality control, assembly, and filtering

An overview of the analysis pipeline is shown in Fig. S16 Quality of raw reads was assessed using FastQC v0.11.9 (124) by examining average quality scores across read sequences to remove samples with average phred scores <20. Assemblies were generated with Shovill v1.1.0 (125) (https://github.com/tseemann/shovill) at default settings, which uses the assembler Spades v3.14 (126) at its core with pre and post-processing steps to refine the final assemblies. The resulting contiguous sequences were assessed for quality using both QUAST v5.2.0 (127) and CheckM v1.2.2 (128). We filtered assemblies using (*i*) CheckM completeness >98%, (*ii*) CheckM contamination <5%, and (*iii*) average genome coverage >10X to retain isolates with high-quality assemblies in the collection. Anomalies in other metrics such as N50, average genome size, and deviation of predicted genes to genome size were negligible once the above three metrics were accounted for.

#### Genome classification and phylogenetic reconstruction

The resulting genome assemblies were taxonomically classified against the Genome Taxonomy Database (GTDB) (Release 07-RS207) (129) using the taxonomic classifier GTDB-Tk v2.1.1 (130). To reconstruct a phylogeny encompassing the diversity of the collection, we first acquired species level representatives using dRep v3.0.0 (131). Within dRep, clustering was conducted with a primary threshold of 90% using MASH v1.1.1 (132), followed by a secondary clustering step using a 95% average nucleotide identity (ANI) threshold with FastANI v1.33 (133).

Ribosomal RNA sequences within each of the species level representatives were predicted using barrnap v0.9 (https://github.com/tseemann/barrnap). Barrnap (134) predicts rRNA sequences using HMMER3 v3.3.2 (135), with models built using the SILVA and Rfam databases. Predicted 16S rRNA sequences were then extracted and aligned using MUSCLE v5.1.0 (136) with default settings. The resulting alignment was trimmed using TrimAL v1.4.1 (137), with a heuristic method optimised for maximum likelihood tree reconstruction (--automated1). The refined alignment was used to reconstruct the phylogenetic tree using IQ-TREE2 v2.0.6 (63) with 1000 ultrafast bootstraps and a TIM3+F+R6 model predicted using ModelFinder (42). The tree was displayed using a 16S rRNA sequence from the *Synergistales* order, *Synergistes jonesii* (GCF_000712295.1), as a root (138). The resulting phylogeny was annotated using the GTDB-Tk taxonomic classifications and the number of isolates per species.

#### Virulence factors

Virulence factors were predicted using ABRicate (139,140) v1.0.1 (https://github.com/tseemann/ABRicate). The default virulence factor database included in ABRicate was replaced with DNA sequences from the full dataset to include all genes related to known and predicted virulence factors (141), (http://www.mgc.ac.cn/VFs/download.html). All gene hits were combined into a single report using the--summary command within ABRicate. Metadata of this database was also examined to describe patterns and biases related to the bacterial origin of these genes (Fig. S6).

#### MosAIC isolates in phylogenetic context

We explored the phylogenetic relationships of three mosquito derived taxa in the collection: *Enterobacter*, *Serratia* and *El. anophelis*. These have defined population structures (60–62); are well represented in the collection and are common mosquito symbionts (21,34). Genomes as described in previous studies (60–62) were retrieved from GenBank (Table S5). Given the limited number of *Enterobacter asburiae* genomes in (61), we included additional *En. asburiae* genomes from the GTDB, filtered by CheckM >98% completeness <5% contamination and existence in a published study (Table S5). Both MosAIC and published genomes were (re)annotated with Bakta v1.7 (142) to ensure consistency. For each of the three datasets, Panaroo v1.3.2 (64) was used to retrieve core gene alignments. Here, all genes present in at least 95% of samples were aligned using MAFFT with clean mode set to moderate, a threshold of 70% identity, and removal of invalid genes such as those with a premature stop codon (--threshold 0.95 –clean-mode moderate-f 0.7-a core –aligner mafft –core_threshold 0.95 –remove-invalid-gene). Variant sites containing single nucleotide polymorphisms (SNPs) were extracted from the core gene alignments using SNP-Sites v2.5.1 (65), allowing for gaps, and the resulting core SNP alignments were used to reconstruct the phylogenies, placing MosAIC isolates in the larger population structures. Each tree was built using 1000 bootstraps with IQ-TREE2 v2.0.6 (63) with the highest scoring models determined using ModelFinder (143); TIM2e+ASC+R4 for *Serratia*, consistent with (62), GTR+F+I+I+R4 for *Enterobacter* and SYM+R3 for *Elizabethkingia*. The *Serratia* phylogenetic tree display was rooted between *Serratia fonticola* and the remaining *Serratia* species, as defined by the original population structure (62); *Enterobacter* and *Elizabethkingia* displays were rooted using the outgroups *Klebsiella aerogenes* and *Chryseobacterium bovis*, respectively.

#### Pangenome analyses

Clusters for the three datasets were defined using PopPUNK v2.6.0 (66), which uses pairwise nucleotide k-mer comparisons to determine shared sequence and gene content and therefore clusters assemblies according to core and accessory gene content. For *En. asburiae* and *El. anophelis*, a Bayesian Gaussian Mixture Model was used to build a network to define clusters. For *S. marcescens*, a HDBSCAN model was used to reflect the larger number of isolates in the phylogeny as recommended by the authors (66). The subsequent refinement step was run using “--fit-model refine” for all predicted clusters to improve overall network scores.

Using PopPUNK-defined lineages and the Panaroo gene presence-absence matrix, we performed population-structure aware gene classifications using the *Twilight* package (67). We set the minimum lineage cluster size to one to account for singleton lineages and core and rare thresholds were set to 0.95 and 0.15, respectively. Gene accumulation curves were generated using the specaccum function within the *vegan* package (144).

All data analysis was conducted in *R* v4.2.2 (145) using the following packages: *tidyverse* v1.3.2 (146), *ggtree* v3.16 (147), *ggtreeExtra* v3.17 (148), *treeio* v3.17 (149), *APE* v5.71 (150), *janitor* (https://github.com/sfirke/janitor) and *phytools* v1.5 (151).

## Data availability

Raw Illumina reads are available in the NCBI Sequence Read Archive (https://www.ncbi.nlm.nih.gov/sra) under BioProject ID PRJNA1023190. Scripts used for analysis and figure generation are available in MosAIC’s GitHub Repository (https://github.com/MosAIC-Collection/MosAIC_V1). All other relevant data supporting the findings of this study, including isolate-specific metadata, are available within the article and its supplementary information.

## Supporting information

File S1 Consortium Author List

## Acknowledgements

We thank Lyric Bartholomay and Kathy Vaccaro (UW-Madison Department of Pathobiological Sciences) for assistance with maintaining the *Ae. aegypti*, *Ae. triseriatus*, *Cx.* pipiens, and *Tx. amboinensis* colonies used to generate material for isolations made by the 2022 UW-Madison Capstone in Microbiology Students included in this study. We also thank Aldo Arellano, Andrew Sommer, and Serena Zhao for assistance with maintenance of internal collections used by personnel in Kerri Coon’s laboratory in the UW-Madison Department of Bacteriology.

## Author contributions

**Conceptualization** Ideas; formulation or evolution of overarching research goals and aims. *GLH, KLC, EH*

**Data Curation** Management activities to annotate (produce metadata), scrub data and maintain research data (including software code, where it is necessary for interpreting the data itself) for initial use and later reuse. *AF, LEB, HLN, KLC, EH*

**Formal Analysis** Application of statistical, mathematical, computational, or other formal techniques to analyze or synthesize study data. *AF, LEB, HLN*

**Funding Acquisition** Acquisition of the financial support for the project leading to this publication. *GLH, KLC, EH*

**Investigation** Conducting a research and investigation process, specifically performing the experiments, or data/evidence collection. *AF, LEB, HLN, MMM, JAL, VD, AFH*

**Methodology** Development or design of methodology; creation of models *AF, LEB, HLN, KLC, EH*

**Project Administration** Management and coordination responsibility for the research activity planning and execution. *GLH, KLC, EH*

**Resources** Provision of study materials, reagents, materials, patients, laboratory samples, animals, instrumentation, computing resources, or other analysis tools. *DEB, EPC, MH, MJL, DJL, EM, CVM, MP, SMS, BS, JX, 2022 UW-Madison Capstone in Microbiology Students, TDP, MRR, GLH, KLC, EH*

**Software** Programming, software development; designing computer programs; implementation of the computer code and supporting algorithms; testing of existing code components. *AF, LEB, EH*

**Supervision** Oversight and leadership responsibility for the research activity planning and execution, including mentorship external to the core team. *LEB, HLN, TDP, MRR, GLH, KLC, EH*

**Validation** Verification, whether as a part of the activity or separate, of the overall replication/reproducibility of results/experiments and other research outputs. *AF, LEB, HLN, MMM, JAL, VD, AFH, KLC, EH*

**Visualization** Preparation, creation and/or presentation of the published work, specifically visualization/data presentation. *AF*

**Writing – Original Draft Preparation** Creation and/or presentation of the published work, specifically writing the initial draft (including substantive translation). *AF, LEB*

**Writing – Review & Editing** Preparation, creation and/or presentation of the published work by those from the original research group, specifically critical review, commentary or revision – including pre-or post-publication stages. *AF, LEB, HLN, GLH, KLC, EH*

*All authors read and approved the final version of the manuscript*.

## Supporting information

**Table S1 |** Curated file containing all isolate-associated metadata, select genome-quality assurance metrics, GTDB classifications, and genome accession numbers. unk = unknown, NA = not applicable.

**Table S2 |** Complete taxonomic classification of all MosAIC isolates, assigned via GTDB-Tk.

**Table S3 |** Virulence factor genes identified in MosAIC isolates using VFDB.

**Table S4 |** Twilight gene classifications for *E. asburiae, E. anophelis* and *S. marcescens* isolates from MosAIC.

**Table S5 |** Metadata for previously sequenced (*i.e.*, external to MosAIC) *Enterobacter*, *Serratia*, and *Elizabethkingia* genomes retrieved from GenBank and manually curated from the GTDB. **File S1 |** Consortium author list.

**Fig. S1 | Distribution of raw sequencing counts.** Histogram showing the size distribution of raw sequencing reads used to assemble MosAIC genomes. The *x*-axis shows the number of reads per isolate, while the *y-*axis shows the number of isolates with a specific read count.

**Fig. S2 | Correlation between number of predicted genes and genome size across MosAIC isolates.** Scatterplot showing the relationship between genome size (*x*-axis) and number of predicted genes (*y*-axis) for each isolate. Each point represents a high-quality genome assembly (>CheckM completeness 98%, <CheckM contamination 5%, >10X read coverage). Line fitted using a linear model in R. Mbp = megabase pairs.

**Fig. S3 | Twenty-five isolates within MosAIC share <95% ANI to a reference genome in the GTDB.** The *x*-axis of the bar chart shows the number of isolates assigned to a reference genome with a given genus assigned taxonomy in the GTDB (*y*-axis). “JAATFS01” is a strain identifier placeholder used by GTDB-Tk when no binomially named representative genome is present in the GTDB.

**Fig. S4 | Bacterial family distribution of MosAIC isolates assigned to different metadata categories.** The *x*-axis of each bar chart shows the number of isolates assigned to a reference genome with a given family assigned taxonomy in the GTDB (*y*-axis). Charts are faceted by metadata category as follows: (A) female_feeding_status, (B) lab_field_derived, (C) mosquito_sex, and (D) mosquito_tissue. Metadata category names and definitions follow those presented in Table S1.

**Fig. S5 | Bacterial genus distribution of MosAIC isolates assigned to different metadata categories.** The *x*-axis of each bar chart shows the number of isolates assigned to a reference genome with a given genus assigned taxonomy in the GTDB (*y*-axis). Charts are faceted by metadata category as follows: (A) lab_field_derived, and (B) mosquito_sex. Metadata category names and definitions follow those presented in Supplementary Table 1. Only isolates assigned to genera within the *Enterobacteriaceae* were included in the analysis.

**Fig. S6 | Taxonomic representation across the VFDB.** Charts are faceted by bacterial class, with the *x*-axis of each chart showing the number of isolates assigned to a given genus on the *y*-axis in which at least one virulence factor gene was identified.

**Fig. S7 | *Elizabethkingia anophelis* Phylogeny with tip labels.** Tip labels of *Elizabethkingia anophelis* shown at each node. Green bar denotes mosquito-associated samples from MosAIC.

**Fig. S8 | *Serratia* Phylogeny with tip labels.** Tip labels of *Serratia* are shown at each node.

**Fig. S9 | *Enterobacter* Phylogeny with tip labels.** Tip labels of *Enterobacter* are shown at each node.

**Fig. S10 | Gene accumulation curve.** Pangenome gene accumulation curve for *En. asburiae, El. anophelis* and *S. marcescens* isolates from MosAIC.

**Fig. S11 | *Enterobacter asburiae* phylogeny overlaid with PopPUNK defined genome clusters.** Tips denote PopPUNK cluster. Green highlight denotes mosquito-associated lineages containing MosAIC isolates.

**Fig. S12 | *Serratia marcescens* phylogeny overlaid with PopPUNK defined genome clusters.** Tips denote PopPUNK cluster. Green highlight denotes mosquito-associated lineages containing MosAIC isolates.

**Fig. S13 | *Elizabethkingia anophelis* phylogeny overlaid with PopPUNK defined genome clusters.** Tips denote PopPUNK cluster. Green highlight denotes mosquito-associated lineages containing MosAIC isolates.

**Fig. S14 | Annotations of lineage-specific core genes identified from mosquito-associated lineages.** Panels summarize annotations for one of three focal species (*Elizabethkingia anophelis, left*; *Serratia marcescens, centre*; or *Enterobacter asburiae, right*), with the *x*-axis of each panel denoting the internal identifier of individual isolates assigned to each species as presented in Table S1. Tiles denote the number of identified annotations corresponding to a given functional category on the *y*-axis, following a gradient from dark blue (few) to light blue (many). White tiles denote categories for which zero annotations were identified in each isolate.

**Fig. S15 | Shared-core gene annotations among mosquito-associated lineages.** Tiles denote the number of identified annotations corresponding to a given functional category on the *y*-axis that were shared between a given species pair on the *x*-axis, following a gradient from dark blue (few) to light blue (many). White tiles denote categories for which zero shared annotations were identified in each species pair.

**Fig. S16 | Analysis workflow.** Flowchart describing (A) the assembly of MosAIC genomes and (B) population genomic analyses.

## References

1. Coon KL, Vogel KJ, Brown MR, Strand MR. Mosquitoes rely on their gut microbiota for development. Mol Ecol. 2014 Jun;23(11):2727–39.

2. Sharma A, Dhayal D, Singh OP, Adak T, Bhatnagar RK. Gut microbes influence fitness and malaria transmission potential of Asian malaria vector Anopheles stephensi. Acta Trop. 2013 Oct;128(1):41–7.

3. Hegde S, Rasgon JL, Hughes GL. The microbiome modulates arbovirus transmission in mosquitoes. Curr Opin Virol. 2015 Dec;15:97–102.

4. Wang J, Gao L, Aksoy S. Microbiota in disease-transmitting vectors. Nat Rev Microbiol. 2023 May 22;21(9):604–18.

5. Cansado-Utrilla C, Zhao SY, McCall PJ, Coon KL, Hughes GL. The microbiome and mosquito vectorial capacity: rich potential for discovery and translation. Microbiome. 2021 Dec;9(1):111.

6. Wang X, Ashraf U, Chen H, Cao S, Ye J. Biological determinants perpetuating the transmission dynamics of mosquito-borne flaviviruses. Emerg Microbes Infect. 2023 Dec 8;12(2):2212812.

7. Ferreira QR, Lemos FFB, Moura MN, Nascimento JODS, Novaes AF, Barcelos IS, et al. Role of the Microbiome in Aedes spp. Vector Competence: What Do We Know? Viruses. 2023 Mar 17;15(3):779.

8. Caragata EP, Short SM. Vector microbiota and immunity: modulating arthropod susceptibility to vertebrate pathogens. Curr Opin Insect Sci. 2022 Apr;50:100875.

9. Caragata EP, Tikhe CV, Dimopoulos G. Curious entanglements: interactions between mosquitoes, their microbiota, and arboviruses. Curr Opin Virol. 2019 Aug;37:26–36.

10. Moyes CL, Vontas J, Martins AJ, Ng LC, Koou SY, Dusfour I, et al. Contemporary status of insecticide resistance in the major Aedes vectors of arboviruses infecting humans. Sinnis P, editor. PLoS Negl Trop Dis. 2017 Jul 20;11(7):e0005625.

11. Ranson H, Lissenden N. Insecticide Resistance in African Anopheles Mosquitoes: A Worsening Situation that Needs Urgent Action to Maintain Malaria Control. Trends Parasitol. 2016 Mar;32(3):187–96.

12. Hegde S, Khanipov K, Albayrak L, Golovko G, Pimenova M, Saldaña MA, et al. Microbiome Interaction Networks and Community Structure From Laboratory-Reared and Field-Collected Aedes aegypti, Aedes albopictus, and Culex quinquefasciatus Mosquito Vectors. Front Microbiol. 2018 Sep 10;9:2160.

13. Saab SA, Dohna H zu, Nilsson LKJ, Onorati P, Nakhleh J, Terenius O, et al. The environment and species affect gut bacteria composition in laboratory co-cultured Anopheles gambiae and Aedes albopictus mosquitoes. Sci Rep. 2020 Dec;10(1):3352.

14. Pérez-Ramos DW, Ramos MM, Payne KC, Giordano BV, Caragata EP. Collection Time, Location, and Mosquito Species Have Distinct Impacts on the Mosquito Microbiota. Front Trop Dis. 2022 May 11;3:896289.

15. Seabourn P, Spafford H, Yoneishi N, Medeiros M. The Aedes albopictus (Diptera: Culicidae) microbiome varies spatially and with Ascogregarine infection. Pimenta PFP, editor. PLoS Negl Trop Dis. 2020 Aug 19;14(8):e0008615.

16. Accoti A, Quek S, Vulcan J, Cansado-Utrilla C, Anderson ER, Alsing J, et al. Microbiome variability of mosquito lines is consistent over time and across environments [Internet]. bioRxiv; 2023. Available from: http://biorxiv.org/lookup/doi/10.1101/2023.04.17.537119

17. Duguma D, Hall MW, Rugman-Jones P, Stouthamer R, Terenius O, Neufeld JD, et al. Developmental succession of the microbiome of Culex mosquitoes. BMC Microbiol. 2015 Dec;15(1):140.

18. Wang X, Liu T, Wu Y, Zhong D, Zhou G, Su X, et al. Bacterial microbiota assemblage in *Aedes albopictus* mosquitoes and its impacts on larval development. Mol Ecol. 2018 Jul;27(14):2972–85.

19. Minard G, Mavingui P, Moro CV. Diversity and function of bacterial microbiota in the mosquito holobiont. Parasit Vectors. 2013 Dec;6(1):146.

20. Zheng R, Wang Q, Wu R, Paradkar PN, Hoffmann AA, Wang GH. Holobiont perspectives on tripartite interactions among microbiota, mosquitoes, and pathogens. ISME J. 2023 May 25;17:1143–52.

21. Scolari F, Casiraghi M, Bonizzoni M. Aedes spp. and Their Microbiota: A Review. Front Microbiol. 2019 Sep 4;10:2036.

22. Bharti R, Grimm DG. Current challenges and best-practice protocols for microbiome analysis. Brief Bioinform. 2021 Jan 18;22(1):178–93.

23. Kozlova EV, Hegde S, Roundy CM, Golovko G, Saldaña MA, Hart CE, et al. Microbial interactions in the mosquito gut determine Serratia colonization and blood-feeding propensity. ISME J. 2021 Jan;15(1):93–108.

24. Stathopoulos S, Neafsey DE, Lawniczak MKN, Muskavitch MAT, Christophides GK. Genetic Dissection of Anopheles gambiae Gut Epithelial Responses to Serratia marcescens. Schneider DS, editor. PLoS Pathog. 2014 Mar 6;10(3):e1003897.

25. Mitraka E, Stathopoulos S, Siden-Kiamos I, Christophides GK, Louis C. *Asaia* accelerates larval development of *Anopheles gambiae*. Pathog Glob Health. 2013 Sep;107(6):305–11.

26. Chouaia B, Rossi P, Epis S, Mosca M, Ricci I, Damiani C, et al. Delayed larval development in Anopheles mosquitoes deprived of Asaia bacterial symbionts. BMC Microbiol. 2012 Dec;12(S1):S2.

27. Cappelli A, Damiani C, Mancini MV, Valzano M, Rossi P, Serrao A, et al. Asaia Activates Immune Genes in Mosquito Eliciting an Anti-Plasmodium Response: Implications in Malaria Control. Front Genet. 2019 Sep 25;10:836.

28. Ezemuoka LC, Akorli EA, Aboagye-Antwi F, Akorli J. Mosquito midgut Enterobacter cloacae and Serratia marcescens affect the fitness of adult female Anopheles gambiae s.l. Terenius O, editor. PLOS ONE. 2020 Sep 18;15(9):e0238931.

29. Jupatanakul N. Serratia marcescens secretes proteases and chitinases with larvicidal activity against Anopheles dirus. Acta Trop. 2020 Aug;212(1):45.

30. Wilke ABB, Marrelli MT. Paratransgenesis: a promising new strategy for mosquito vector control. Parasit Vectors. 2015 Dec;8(1):342.

31. Wang S, Jacobs-Lorena M. Paratransgenesis Applications. In: Arthropod Vector: Controller of Disease Transmission, Volume 1 [Internet]. Elsevier; 2017. p. 219–34. Available from: https://linkinghub.elsevier.com/retrieve/pii/B9780128053508000131

32. Elston KM, Leonard SP, Geng P, Bialik SB, Robinson E, Barrick JE. Engineering insects from the endosymbiont out. Trends Microbiol. 2022 Jan;30(1):79–96.

33. Huang W, Rodrigues J, Bilgo E, Tormo JR, Challenger JD, Cozar-Gallardo CD, et al. Delftia tsuruhatensis TC1 symbiont suppresses malaria transmission by anopheline mosquitoes. Science. 2023;381(6657):533–40.

34. Steven B, Hyde J, LaReau JC, Brackney DE. The Axenic and Gnotobiotic Mosquito: Emerging Models for Microbiome Host Interactions. Front Microbiol. 2021 Jul 12;12:714222.

35. Romoli O, Schönbeck JC, Hapfelmeier S, Gendrin M. Production of germ-free mosquitoes via transient colonisation allows stage-specific investigation of host-microbiota interactions. Nat Commun [Internet]. 2021 Feb 11;12(942). Available from: https://www.nature.com/articles/s41467-021-21195-3

36. Wu J, Wang Q, Wang D, Wong ACN, Wang GH. Axenic and gnotobiotic insect technologies in research on host–microbiota interactions. Trends Microbiol. 2023 Aug;31(8):858–71.

37. Zhao SY, Hughes GL, Coon KL. A cryopreservation method to recover laboratory-and field-derived bacterial communities from mosquito larval habitats. Bartholomay LC, editor. PLoS Negl Trop Dis. 2023 Apr 5;17(4):e0011234.

38. Coon KL, Hegde S, Hughes GL. Inter-species microbiota transplantation recapitulates microbial acquisition and persistence in mosquitoes. Microbiome [Internet]. 2022 Apr 11;10(58). Available from: https://microbiomejournal.biomedcentral.com/articles/10.1186/s40168-022-01256-5

39. Hegde S, Nilyanimit P, Kozlova E, Anderson ER, Narra HP, Sahni SK, et al. CRISPR/Cas9-mediated gene deletion of the ompA gene in symbiotic Cedecea neteri impairs biofilm formation and reduces gut colonization of Aedes aegypti mosquitoes. Bonizzoni M, editor. PLoS Negl Trop Dis. 2019 Dec 2;13(12):e0007883.

40. Hegde S, Brettell LE, Quek S, Etebari K, Saldaña MA, Asgari S, et al. Aedes aegypti gut transcriptomes respond differently to microbiome transplants from field-caught or laboratory-reared mosquitoes. 2023 Mar 16; Available from: http://biorxiv.org/lookup/doi/10.1101/2023.03.16.532926

41. Dickson LB, Jiolle D, Minard G, Moltini-Conclois I, Volant S, Ghozlane A, et al. Carryover effects of larval exposure to different environmental bacteria drive adult trait variation in a mosquito vector. Sci Adv. 2017 Aug;3(8):e1700585.

42. Dennison NJ, Saraiva RG, Cirimotich CM, Mlambo G, Mongodin EF, Dimopoulos G. Functional genomic analyses of Enterobacter, Anopheles and Plasmodium reciprocal interactions that impact vector competence. Malar J. 2016 Dec;15(1):425.

43. Tchioffo MT, Boissière A, Churcher TS, Abate L, Gimonneau G, Nsango SE, et al. Modulation of Malaria Infection in Anopheles gambiae Mosquitoes Exposed to Natural Midgut Bacteria. Michel K, editor. PLoS ONE. 2013 Dec 6;8(12):e81663.

44. Ganley JG, Pandey A, Sylvester K, Lu KY, Toro-Moreno M, Rütschlin S, et al. A Systematic Analysis of Mosquito-Microbiome Biosynthetic Gene Clusters Reveals Antimalarial Siderophores that Reduce Mosquito Reproduction Capacity. Cell Chem Biol. 2020 Jul;27(7):817–826.e5.

45. Strand MR. Composition and functional roles of the gut microbiota in mosquitoes. Curr Opin Insect Sci. 2018 Aug;28:59–65.

46. Coon KL, Valzania L, McKinney DA, Vogel KJ, Brown MR, Strand MR. Bacteria-mediated hypoxia functions as a signal for mosquito development. Proc Natl Acad Sci. 2017 Jul 3;114(27):E5362–9.

47. Coon KL, Brown MR, Strand MR. Mosquitoes host communities of bacteria that are essential for development but vary greatly between local habitats. Mol Ecol. 2016 Nov;25(22):5806–26.

48. Valzania L, Martinson VG, Harrison RE, Boyd BM, Coon KL, Brown MR, et al. Both living bacteria and eukaryotes in the mosquito gut promote growth of larvae. Bartholomay LC, editor. PLoS Negl Trop Dis. 2018 Jul 6;12(7):e0006638.

49. Girard M, Martin E, Vallon L, Raquin V, Bellet C, Rozier Y, et al. Microorganisms Associated with Mosquito Oviposition Sites: Implications for Habitat Selection and Insect Life Histories. Microorganisms. 2021 Jul 26;9(8):1589.

50. Barnard K, Jeanrenaud ACSN, Brooke BD, Oliver SV. The contribution of gut bacteria to insecticide resistance and the life histories of the major malaria vector Anopheles arabiensis (Diptera: Culicidae). Sci Rep. 2019 Jun 24;9(1):9117.

51. Schmidt K, Engel P. Mechanisms underlying gut microbiota–host interactions in insects. J Exp Biol. 2021 Jan 15;224(2):jeb207696.

52. Chen S, Blom J, Walker ED. Genomic, Physiologic, and Symbiotic Characterization of Serratia marcescens Strains Isolated from the Mosquito Anopheles stephensi. Front Microbiol. 2017 Aug 10;8:1483.

53. Watson M. New insights from 33,813 publicly available metagenome-assembled-genomes (MAGs) assembled from the rumen microbiome. 2021 Apr 2; Available from: http://biorxiv.org/lookup/doi/10.1101/2021.04.02.438222

54. Glendinning L, Stewart RD, Pallen MJ, Watson KA, Watson M. Assembly of hundreds of novel bacterial genomes from the chicken caecum. Genome Biol. 2020 Dec;21(1):34.

55. Stewart RD, Auffret MD, Warr A, Wiser AH, Press MO, Langford KW, et al. Assembly of 913 microbial genomes from metagenomic sequencing of the cow rumen. Nat Commun. 2018 Dec;9(1):870.

56. Wilkinson T, Korir D, Ogugo M, Stewart RD, Watson M, Paxton E, et al. 1200 high-quality metagenome-assembled genomes from the rumen of African cattle and their relevance in the context of sub-optimal feeding. Genome Biol. 2020 Dec;21(1):229.

57. Almeida A, Mitchell AL, Boland M, Forster SC, Gloor GB, Tarkowska A, et al. A new genomic blueprint of the human gut microbiota. Nature. 2019 Apr 25;568(7753):499–504.

58. Land M, Hauser L, Jun SR, Nookaew I, Leuze MR, Ahn TH, et al. Insights from 20 years of bacterial genome sequencing. Funct Integr Genomics. 2015 Mar;15(2):141–61.

59. Olm MR, Crits-Christoph A, Diamond S, Lavy A, Matheus Carnevali PB, Banfield JF. Consistent Metagenome-Derived Metrics Verify and Delineate Bacterial Species Boundaries. Woyke T, editor. mSystems. 2020 Feb 11;5(1):e00731-19.

60. Hu S, Xu H, Meng X, Bai X, Xu J, Ji J, et al. Population genomics of emerging *Elizabethkingia anophelis* pathogens reveals potential outbreak and rapid global dissemination. Emerg Microbes Infect. 2022 Dec 31;11(1):2590–9.

61. Chavda KD, Chen L, Fouts DE, Sutton G, Brinkac L, Jenkins SG, et al. Comprehensive Genome Analysis of Carbapenemase-Producing *Enterobacter* spp.: New Insights into Phylogeny, Population Structure, and Resistance Mechanisms. Jacoby GA, editor. mBio. 2016 Dec 30;7(6):e02093-16.

62. Williams DJ, Grimont PAD, Cazares A, Grimont F, Ageron E, Pettigrew KA, et al. The genus Serratia revisited by genomics. Nat Commun. 2022 Sep 3;13(1):5195.

63. Minh BQ, Schmidt HA, Chernomor O, Schrempf D, Woodhams MD, Von Haeseler A, et al. IQ-TREE 2: New Models and Efficient Methods for Phylogenetic Inference in the Genomic Era. Teeling E, editor. Mol Biol Evol. 2020 May 1;37(5):1530–4.

64. Tonkin-Hill G, MacAlasdair N, Ruis C, Weimann A, Horesh G, Lees JA, et al. Producing polished prokaryotic pangenomes with the Panaroo pipeline. Genome Biol. 2020 Dec;21(1):180.

65. Page AJ, Taylor B, Delaney AJ, Soares J, Seemann T, Keane A, et al. SNP-sites: rapid efficient extraction of SNPs from multi-FASTA alignments. Microb Genomics.

66. Lees JA, Harris SR, Tonkin-Hill G, Gladstone RA, Lo SW, Weiser JN, et al. Fast and flexible bacterial genomic epidemiology with PopPUNK. Genome Res. 2019 Feb;29(2):304–16.

67. Horesh G, Taylor-Brown A, McGimpsey S, Lassalle F, Corander J, Heinz E, et al. Different evolutionary trends form the twilight zone of the bacterial pan-genome. Microb Genomics [Internet]. 2021 Sep 24;7(9). Available from: https://www.microbiologyresearch.org/content/journal/mgen/10.1099/mgen.0.000670

68. Coon KL, Strand MR. Gut Microbiome Assembly and Function in Mosquitoes. In: Population Biology of Vector-Borne Diseases [Internet]. Oxford University Press; 2020. p. 227–44. Available from: https://oxford.universitypressscholarship.com/view/10.1093/oso/9780198853244.001.0001/ oso-9780198853244-chapter-13

69. Dada N, Jupatanakul N, Minard G, Short SM, Akorli J, Villegas LM. Considerations for mosquito microbiome research from the Mosquito Microbiome Consortium. Microbiome. 2021 Dec;9(1):36.

70. Guégan M, Zouache K, Démichel C, Minard G, Tran Van V, Potier P, et al. The mosquito holobiont: fresh insight into mosquito-microbiota interactions. Microbiome. 2018 Dec;6(1):49.

71. Benfey PN, Mitchell-Olds T. From Genotype to Phenotype: Systems Biology Meets Natural Variation. Science. 2008 Apr 25;320(5875):495–7.

72. Muturi EJ, Njoroge TM, Dunlap C, Cáceres CE. Blood meal source and mixed blood-feeding influence gut bacterial community composition in Aedes aegypti. Parasit Vectors. 2021 Dec;14(1):83.

73. Bennett KL, Gómez-Martínez C, Chin Y, Saltonstall K, McMillan WO, Rovira JR, et al. Dynamics and diversity of bacteria associated with the disease vectors Aedes aegypti and Aedes albopictus. Sci Rep. 2019 Dec;9(1):12160.

74. Cimermancic P, Medema MH, Claesen J, Kurita K, Wieland Brown LC, Mavrommatis K, et al. Insights into Secondary Metabolism from a Global Analysis of Prokaryotic Biosynthetic Gene Clusters. Cell. 2014 Jul;158(2):412–21.

75. Whiten SR, Eggleston H, Adelman ZN. Ironing out the Details: Exploring the Role of Iron and Heme in Blood-Sucking Arthropods. Front Physiol. 2018 Jan 17;8:1134.

76. Onyango MG, Lange R, Bialosuknia S, Payne A, Mathias N, Kuo L, et al. Zika virus and temperature modulate Elizabethkingia anophelis in Aedes albopictus. Parasit Vectors. 2021 Dec;14(1):573.

77. Wang Y, Gilbreath TM, Kukutla P, Yan G, Xu J. Dynamic Gut Microbiome across Life History of the Malaria Mosquito Anopheles gambiae in Kenya. Leulier F, editor. PLoS ONE. 2011 Sep 21;6(9):e24767.

78. Muturi EJ, Dunlap C, Ramirez JL, Rooney AP, Kim CH. Host blood meal source has a strong impact on gut microbiota of Aedes aegypti. FEMS Microbiol Ecol [Internet]. 2018 Oct 24;95(1). Available from: https://academic.oup.com/femsec/advance-article/doi/10.1093/femsec/fiy213/5144212

79. Gusmão DS, Santos AV, Marini DC, Bacci M, Berbert-Molina MA, Lemos FJA. Culture-dependent and culture-independent characterization of microorganisms associated with Aedes aegypti (Diptera: Culicidae) (L.) and dynamics of bacterial colonization in the midgut. Acta Trop. 2010 Sep;115(3):275–81.

80. Thapa S, Pant ND, Shrestha R, Gc G, Shrestha B, Pandey BD, et al. Prevalence of dengue and diversity of cultivable bacteria in vector Aedes aegypti (L.) from two dengue endemic districts, Kanchanpur and Parsa of Nepal. J Health Popul Nutr. 2017 Dec;36(1):5.

81. Valiente Moro C, Tran FH, Nantenaina Raharimalala F, Ravelonandro P, Mavingui P. Diversity of culturable bacteria including Pantoea in wild mosquito Aedes albopictus. BMC Microbiol. 2013;13(1):70.

82. Chen S, Johnson BK, Yu T, Nelson BN, Walker ED. Elizabethkingia anophelis: Physiologic and Transcriptomic Responses to Iron Stress. Front Microbiol. 2020 May 7;11:804.

83. Chen S, Bagdasarian M, Walker ED. Elizabethkingia anophelis: Molecular Manipulation and Interactions with Mosquito Hosts. Schottel JL, editor. Appl Environ Microbiol. 2015 Mar 15;81(6):2233–43.

84. Ganley JG, D’Ambrosio HK, Shieh M, Derbyshire ER. Coculturing of Mosquito-Microbiome Bacteria Promotes Heme Degradation in *Elizabethkingia anophelis*. ChemBioChem. 2020 May 4;21(9):1279–84.

85. Kukutla P, Lindberg BG, Pei D, Rayl M, Yu W, Steritz M, et al. Insights from the Genome Annotation of Elizabethkingia anophelis from the Malaria Vector Anopheles gambiae. Tu Z, editor. PLoS ONE. 2014 May 19;9(5):e97715.

86. Díaz-Nieto LM, D’Alessio C, Perotti MA, Berón CM. Culex pipiens Development Is Greatly Influenced by Native Bacteria and Exogenous Yeast. Terenius O, editor. PLOS ONE. 2016 Apr 7;11(4):e0153133.

87. Buck M, Nilsson LKJ, Brunius C, Dabiré RK, Hopkins R, Terenius O. Bacterial associations reveal spatial population dynamics in Anopheles gambiae mosquitoes. Sci Rep. 2016 Mar 10;6(1):22806.

88. Chouaia B, Gaiarsa S, Crotti E, Comandatore F, Degli Esposti M, Ricci I, et al. Acetic Acid Bacteria Genomes Reveal Functional Traits for Adaptation to Life in Insect Guts. Genome Biol Evol. 2014 Apr;6(4):912–20.

89. Crotti E, Rizzi A, Chouaia B, Ricci I, Favia G, Alma A, et al. Acetic Acid Bacteria, Newly Emerging Symbionts of Insects. Appl Environ Microbiol. 2010 Nov;76(21):6963–70.

90. Lagkouvardos I, Pukall R, Abt B, Foesel BU, Meier-Kolthoff JP, Kumar N, et al. The Mouse Intestinal Bacterial Collection (miBC) provides host-specific insight into cultured diversity and functional potential of the gut microbiota. Nat Microbiol. 2016 Aug 8;1(10):16131.

91. Li W, Liang H, Lin X, Hu T, Wu Z, He W, et al. A catalog of bacterial reference genomes from cultivated human oral bacteria. Npj Biofilms Microbiomes. 2023 Jul 3;9(1):45.

92. Alfano N, Tagliapietra V, Rosso F, Manica M, Arnoldi D, Pindo M, et al. Changes in Microbiota Across Developmental Stages of Aedes koreicus, an Invasive Mosquito Vector in Europe: Indications for Microbiota-Based Control Strategies. Front Microbiol. 2019 Dec 10;10:2832.

93. Moll RM, Romoser WS, Modrakowski MC, Moncayo AC, Lerdthusnee K. Meconial Peritrophic Membranes and the Fate of Midgut Bacteria During Mosquito (Diptera: Culicidae) Metamorphosis. J Med Entomol. 2001 Jan 1;38(1):29–32.

94. Muturi EJ, Ramirez JL, Rooney AP, Kim CH. Comparative analysis of gut microbiota of mosquito communities in central Illinois. Kittayapong P, editor. PLoS Negl Trop Dis. 2017 Feb 28;11(2):e0005377.

95. Zouache K, Raharimalala FN, Raquin V, Tran-Van V, Raveloson LHR, Ravelonandro P, et al. Bacterial diversity of field-caught mosquitoes, Aedes albopictus and Aedes aegypti, from different geographic regions of Madagascar: Bacterial communities of wild Aedes mosquito vectors. FEMS Microbiol Ecol. 2011 Mar;75(3):377–89.

96. Boissière A, Tchioffo MT, Bachar D, Abate L, Marie A, Nsango SE, et al. Midgut Microbiota of the Malaria Mosquito Vector Anopheles gambiae and Interactions with Plasmodium falciparum Infection. Vernick KD, editor. PLoS Pathog. 2012 May 31;8(5):e1002742.

97. Osei-Poku J, Mbogo CM, Palmer WJ, Jiggins FM. Deep sequencing reveals extensive variation in the gut microbiota of wild mosquitoes from Kenya. Mol Ecol. 2012 Oct;21(20):5138–50.

98. Gimonneau G, Tchioffo MT, Abate L, Boissière A, Awono-Ambéné PH, Nsango SE, et al. Composition of Anopheles coluzzii and Anopheles gambiae microbiota from larval to adult stages. Infect Genet Evol. 2014 Dec;28:715–24.

99. Staudacher H, Kaltenpoth M, Breeuwer JAJ, Menken SBJ, Heckel DG, Groot AT. Variability of Bacterial Communities in the Moth Heliothis virescens Indicates Transient Association with the Host. Abdo Z, editor. PLOS ONE. 2016 May 3;11(5):e0154514.

100. Paniagua Voirol LR, Frago E, Kaltenpoth M, Hilker M, Fatouros NE. Bacterial Symbionts in Lepidoptera: Their Diversity, Transmission, and Impact on the Host. Front Microbiol. 2018 Mar 27;9:556.

101. Ellegaard KM, Engel P. Genomic diversity landscape of the honey bee gut microbiota. Nat Commun. 2019 Jan 25;10(1):446.

102. Brune A, Dietrich C. The Gut Microbiota of Termites: Digesting the Diversity in the Light of Ecology and Evolution. Annu Rev Microbiol. 2015 Oct 15;69(1):145–66.

103. Sheppard SK, Guttman DS, Fitzgerald JR. Population genomics of bacterial host adaptation. Nat Rev Genet. 2018 Sep;19(9):549–65.

104. Winans NJ, Walter A, Chouaia B, Chaston JM, Douglas AE, Newell PD. A genomic investigation of ecological differentiation between free-living and *Drosophila*-associated bacteria. Mol Ecol. 2017 Sep;26(17):4536–50.

105. Zhou G, Kohlhepp P, Geiser D, Winzerling JJ. Fate of blood meal iron in mosquitos. J Insect Physiol. 2008;53(11):1169–78.

106. Maya-Maldonado K, Cardoso-Jaime V, González-Olvera G, Osorio B, Recio-Tótoro B, Manrique-Saide P, et al. Mosquito metallomics reveal copper and iron as critical factors for Plasmodium infection. Mireji PO, editor. PLoS Negl Trop Dis. 2021 Jun 23;15(6):e0009509.

107. Benoit JB, Lopez-Martinez G, Patrick KR, Phillips ZP, Krause TB, Denlinger DL. Drinking a hot blood meal elicits a protective heat shock response in mosquitoes. Proc Natl Acad Sci. 2011 May 10;108(19):8026–9.

108. Billker O, Miller AJ, Sinden RE. Determination of mosquito bloodmeal pH *in situ* by ion-selective microelectrode measurement: implications for the regulation of malarial gametogenesis. Parasitology. 2000 Jun;120(6):547–51.

109. Kakani P, Gupta L, Kumar S. Heme-Peroxidase 2, a Peroxinectin-Like Gene, Regulates Bacterial Homeostasis in Anopheles stephensi Midgut. Front Physiol. 2020 Sep 8;11:572340.

110. Burgsdorf I, Handley KM, Bar-Shalom R, Erwin PM, Steindler L. Life at Home and on the Roam: Genomic Adaptions Reflect the Dual Lifestyle of an Intracellular, Facultative Symbiont. Bordenstein S, editor. mSystems. 2019 Aug 27;4(4):e00057-19.

111. Calkins TL, DeLaat A, Piermarini PM. Physiological characterization and regulation of the contractile properties of the mosquito ventral diverticulum (crop). J Insect Physiol. 2017 Nov;103:98–106.

112. Hall-Stoodley L, Costerton JW, Stoodley P. Bacterial biofilms: from the Natural environment to infectious diseases. Nat Rev Microbiol. 2004 Feb;2(2):95–108.

113. Kinosita Y, Kikuchi Y, Mikami N, Nakane D, Nishizaka T. Unforeseen swimming and gliding mode of an insect gut symbiont, Burkholderia sp. RPE64, with wrapping of the flagella around its cell body. ISME J. 2018 Mar;12(3):838–48.

114. Ganesan R, Wierz JC, Kaltenpoth M, Flórez LV. How It All Begins: Bacterial Factors Mediating the Colonization of Invertebrate Hosts by Beneficial Symbionts. Microbiol Mol Biol Rev. 2022 Dec 21;86(4):e00126–21.

115. Foo A, Cerdeira L, Hughes GL, Heinz E. Recovery of Metagenomic Data from the *Aedes aegypti* Microbiome using a Reproducible Snakemake Pipeline: MINUUR. Wellcome Open [Internet]. 2023 May 26;8(131). Available from: https://wellcomeopenresearch.org/articles/8-131

116. Pérez-Cobas AE, Gomez-Valero L, Buchrieser C. Metagenomic approaches in microbial ecology: an update on whole-genome and marker gene sequencing analyses. Microb Genomics [Internet]. 2020 Aug 1;6(8). Available from: https://www.microbiologyresearch.org/content/journal/mgen/10.1099/mgen.0.000409

117. Jiao JY, Liu L, Hua ZS, Fang BZ, Zhou EM, Salam N, et al. Microbial dark matter coming to light: challenges and opportunities. Natl Sci Rev. 2021 Mar 19;8(3):nwaa280.

118. Doolittle WF, Booth A. It’s the song, not the singer: an exploration of holobiosis and evolutionary theory. Biol Philos. 2017 Jan;32(1):5–24.

119. Louca S, Jacques SMS, Pires APF, Leal JS, Srivastava DS, Parfrey LW, et al. High taxonomic variability despite stable functional structure across microbial communities. Nat Ecol Evol. 2016 Dec 5;1(1):0015.

120. Burke C, Steinberg P, Rusch D, Kjelleberg S, Thomas T. Bacterial community assembly based on functional genes rather than species. Proc Natl Acad Sci. 2011 Aug 23;108(34):14288–93.

121. The Human Microbiome Project Consortium. Structure, function and diversity of the healthy human microbiome. Nature. 2012 Jun;486(7402):207–14.

122. Turnbaugh PJ, Hamady M, Yatsunenko T, Cantarel BL, Duncan A, Ley RE, et al. A core gut microbiome in obese and lean twins. Nature. 2009 Jan;457(7228):480–4.

123. Power RA, Parkhill J, De Oliveira T. Microbial genome-wide association studies: lessons from human GWAS. Nat Rev Genet. 2017 Jan;18(1):41–50.

124. Andrew, S A S. FASTQC: A Quality Control Tool for High Throughput Sequence Data [Online] [Internet]. Available from: www.bioinformatics.babraham.ac.uk/projects/fastqc/

125. Rahmani A, Meradi L, Piris R. Molecular characterization of the whole genome in clinical multidrug-resistant strains of Klebsiella pneumoniae. J Infect Dev Ctries. 2023 Jan 31;17(01):59–65.

126. Bankevich A, Nurk S, Antipov D, Gurevich AA, Dvorkin M, Kulikov AS, et al. SPAdes: A New Genome Assembly Algorithm and Its Applications to Single-Cell Sequencing. J Comput Biol. 2012 May;19(5):455–77.

127. Gurevich A, Saveliev V, Vyahhi N, Tesler G. QUAST: quality assessment tool for genome assemblies. Bioinformatics. 2013 Apr 15;29(8):1072–5.

128. Parks DH, Imelfort M, Skennerton CT, Hugenholtz P, Tyson GW. CheckM: assessing the quality of microbial genomes recovered from isolates, single cells, and metagenomes. Genome Res. 2015 Jul;25(7):1043–55.

129. Hugenholtz P, Parks DH. Genome Taxonomy Database r207. Version 1.92. The University of Queensland; 2022.

130. Chaumeil PA, Mussig AJ, Hugenholtz P, Parks DH. GTDB-Tk v2: memory friendly classification with the genome taxonomy database. Borgwardt K, editor. Bioinformatics. 2022 Nov 30;38(23):5315–6.

131. Olm MR, Brown CT, Brooks B, Banfield JF. dRep: a tool for fast and accurate genomic comparisons that enables improved genome recovery from metagenomes through de-replication. ISME J. 2017 Dec;11(12):2864–8.

132. Ondov BD, Treangen TJ, Melsted P, Mallonee AB, Bergman NH, Koren S, et al. Mash: fast genome and metagenome distance estimation using MinHash. Genome Biol. 2016 Dec;17(1):132.

133. Jain C, Rodriguez-R LM, Phillippy AM, Konstantinidis KT, Aluru S. High throughput ANI analysis of 90K prokaryotic genomes reveals clear species boundaries. Nat Commun. 2018 Nov 30;9(1):5114.

134. Liu N, Zhu L, Zhang Z, Huang H, Jiang L. Draft genome sequence of a multidrug-resistant blaOXA-69-producing Acinetobacter baumannii L13 isolated from Tarim River sample in China. J Glob Antimicrob Resist. 2019 Sep;18:145–7.

135. Finn RD, Clements J, Eddy SR. HMMER web server: interactive sequence similarity searching. Nucleic Acids Res. 2011 Jul 1;39:W29–37.

136. Edgar RC. MUSCLE - a multiple sequence alignment method with reduced time and space complexity. BMC Bioinformatics. 2004;5(1):113.

137. Capella-Gutierrez S, Silla-Martinez JM, Gabaldon T. trimAl: a tool for automated alignment trimming in large-scale phylogenetic analyses. Bioinformatics. 2009 Aug 1;25(15):1972–3.

138. Zhu Q, Mai U, Pfeiffer W, Janssen S, Asnicar F, Sanders JG, et al. Phylogenomics of 10,575 genomes reveals evolutionary proximity between domains Bacteria and Archaea. Nat Commun. 2019 Dec 2;10(1):5477.

139. Camargo A, Guerrero-Araya E, Castañeda S, Vega L, Cardenas-Alvarez MX, Rodríguez C, et al. Intra-species diversity of Clostridium perfringens: A diverse genetic repertoire reveals its pathogenic potential. Front Microbiol. 2022 Jul 22;13:952081.

140. Tian C, Xing M, Fu L, Zhao Y, Fan X, Wang S. Emergence of uncommon KL38-OCL6-ST220 carbapenem-resistant Acinetobacter pittii strain, co-producing chromosomal NDM-1 and OXA-820 carbapenemases. Front Cell Infect Microbiol. 2022 Aug 12;12:943735.

141. Liu B, Zheng D, Zhou S, Chen L, Yang J. VFDB 2022: a general classification scheme for bacterial virulence factors. Nucleic Acids Res. 2022 Jan 7;50(D1):D912–7.

142. Schwengers O, Jelonek L, Dieckmann MA, Beyvers S, Blom J, Goesmann A. Bakta: rapid and standardized annotation of bacterial genomes via alignment-free sequence identification. Microb Genomics [Internet]. 2021 Nov 5 [cited 2023 Mar 23];7(11). Available from: https://www.microbiologyresearch.org/content/journal/mgen/10.1099/mgen.0.000685

143. Kalyaanamoorthy S, Minh BQ, Wong TKF, Von Haeseler A, Jermiin LS. ModelFinder: fast model selection for accurate phylogenetic estimates. Nat Methods. 2017 Jun;14(6):587–9.

144. Dixon P. VEGAN, a package of R functions for community ecology. J Veg Sci. 2003 Dec;14(6):927–30.

145. R Core Team. R: A language and environment for statistical computing. R foundation for statistical computing, Vienna, Austria [Internet]. 2021. Available from: https://www.R-project.org/

146. Wickham H, Averick M, Bryan J, Chang W, McGowan L, François R, et al. Welcome to the Tidyverse. J Open Source Softw. 2019 Nov 21;4(43):1686.

147. Yu G, Smith DK, Zhu H, Guan Y, Lam TT. GGTREE: an R package for visualization and annotation of phylogenetic trees with their covariates and other associated data. McInerny G, editor. Methods Ecol Evol. 2017 Jan;8(1):28–36.

148. Xu S, Dai Z, Guo P, Fu X, Liu S, Zhou L, et al. ggtreeExtra: Compact Visualization of Richly Annotated Phylogenetic Data. Tamura K, editor. Mol Biol Evol. 2021 Aug 23;38(9):4039–42.

149. Wang LG, Lam TTY, Xu S, Dai Z, Zhou L, Feng T, et al. Treeio: An R Package for Phylogenetic Tree Input and Output with Richly Annotated and Associated Data. Kumar S, editor. Mol Biol Evol. 2020 Feb 1;37(2):599–603.

150. Paradis E, Schliep K. ape 5.0: an environment for modern phylogenetics and evolutionary analyses in R. Schwartz R, editor. Bioinformatics. 2019 Feb 1;35(3):526–8.

151. Revell LJ. phytools: an R package for phylogenetic comparative biology (and other things): *phytools: R package*. Methods Ecol Evol. 2012 Apr;3(2):217–23.

