## Supplementary material for "Establishment and comparative genomics of a high-quality collection of mosquito-associated bacterial isolates - MosAIC (Mosquito-Associated Isolate Collection)": File S1 Consortium Author List

Supplementary File 1: Consortium Author List

2022 UW-Madison Capstone in Microbiology Students

Consortium authors include all persons enrolled in MICROBIO 551 (Capstone Research Project in Microbiology, 2 cr.) during the Spring 2022 semester that (*i*) were identified by Timothy Paustian (Teaching Professor, Department of Bacteriology) or Michelle Rondon (Teaching Faculty, Department of Bacteriology) as deserving co-authorship credit, and (*ii*) gave consent to have their name published alongside this study. These individuals are listed alphabetically below.

| Shraddha R. Bhide |
| --- |
| Annika H. Borgaonkar |
| Jakob J. Burkett |
| Daniel J. Chacko |
| Nikolas L. Christoffel |
| Joy Chung |
| Andrew J. DeMarco |
| Nicholas P. Durst |
| Pratyusha Emkay |
| Rebecca N. Forman |
| Adalee K. Gill |
| Uma A. Gude |
| Giovanni M. Hanstad |
| Lars H. Johnston |
| Matthew R. Johnston |
| Arman Kamrani |
| Martin Kelty |
| Noah R. Kokko-Ludemann |
| Kennah E. Konrad |
| Michael D. LeClaire |
| Grace Lin |
| Robert C. Londono |
| Jillian M. Lucito |
| Katelyn R. Major |
| Thomas J. Montana |
| John G. Price |
| Sarah A. Sabinash |
| Ann M. Saucedo |
| Allison A. Schopf |
| Anna Schwenn |
| Natalee H. Skoubis |
| Dawson J. Slusser |
| Justin Stensloff |
| Dianne L. Tebbe |
| Addison Vang |
