## Supplementary material for "Establishment and comparative genomics of a high-quality collection of mosquito-associated bacterial isolates - MosAIC (Mosquito-Associated Isolate Collection)": Figures S1 to S16

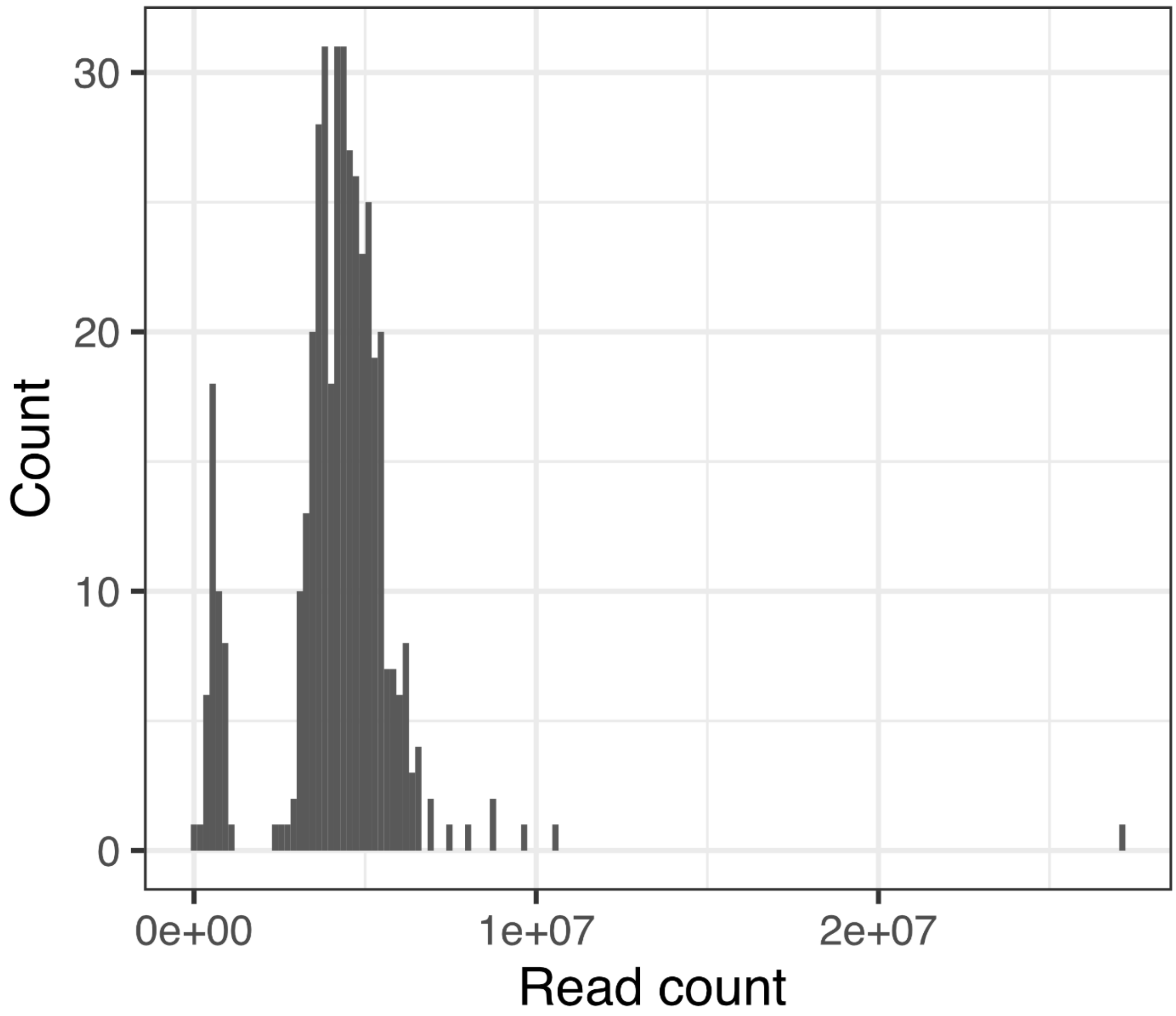


**Fig. S1 | Distribution of raw sequencing counts.** Histogram showing the size distribution of raw sequencing reads used to assemble MosAIC genomes. The *x*-axis shows the number of reads per isolate, while the *y*-axis shows the number of isolates with a specific read count.

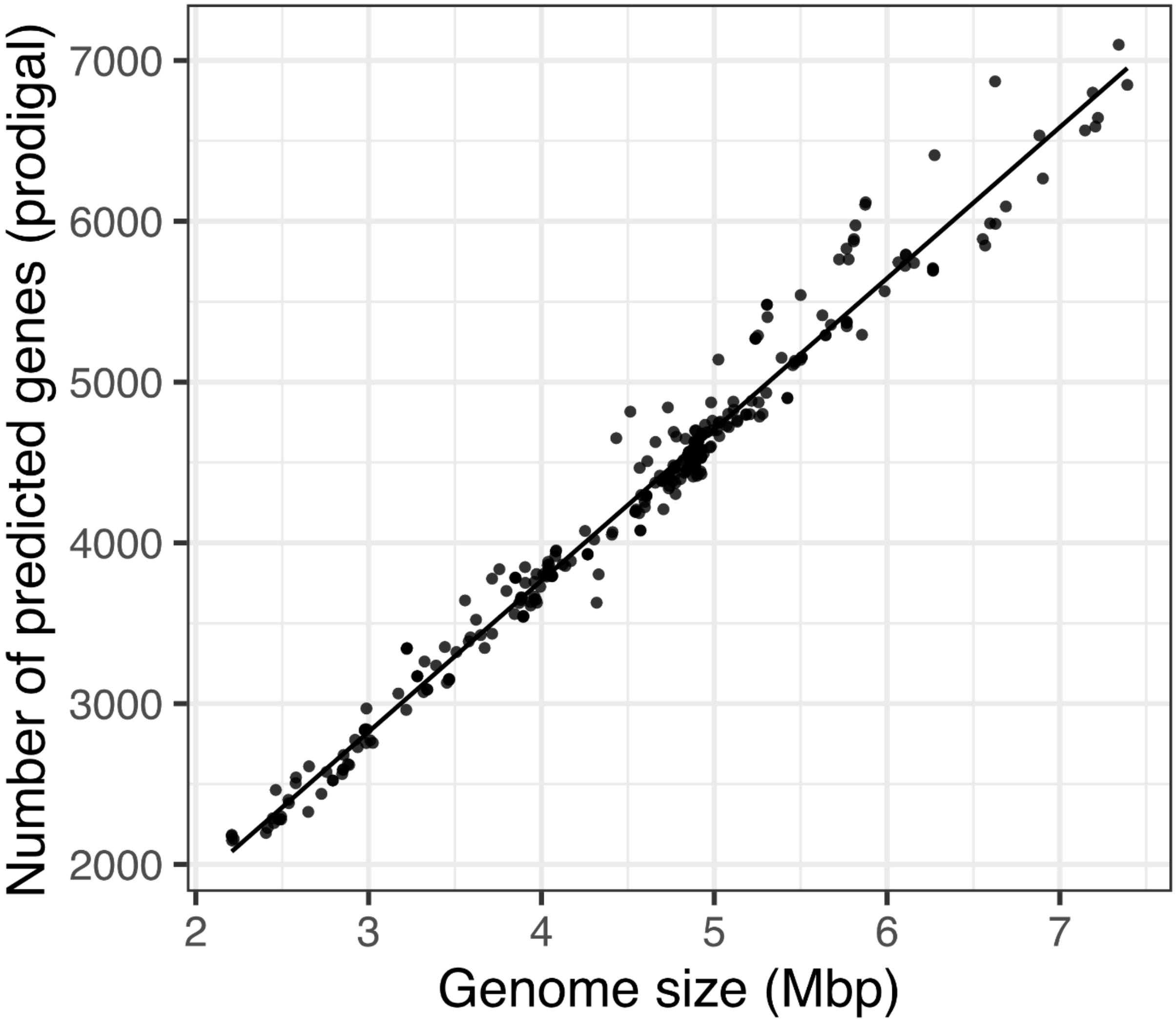


**Fig. S2 | Correlation between number of predicted genes and genome size across MosAIC isolates.** Scatterplot showing the relationship between genome size (*x*-axis) and number of predicted genes (*y*-axis) for each isolate. Each point represents a high-quality genome assembly (>CheckM completeness 98%, <CheckM contamination 5%, >10X read coverage). Line fitted using a linear model in R. Mbp = megabase pairs.

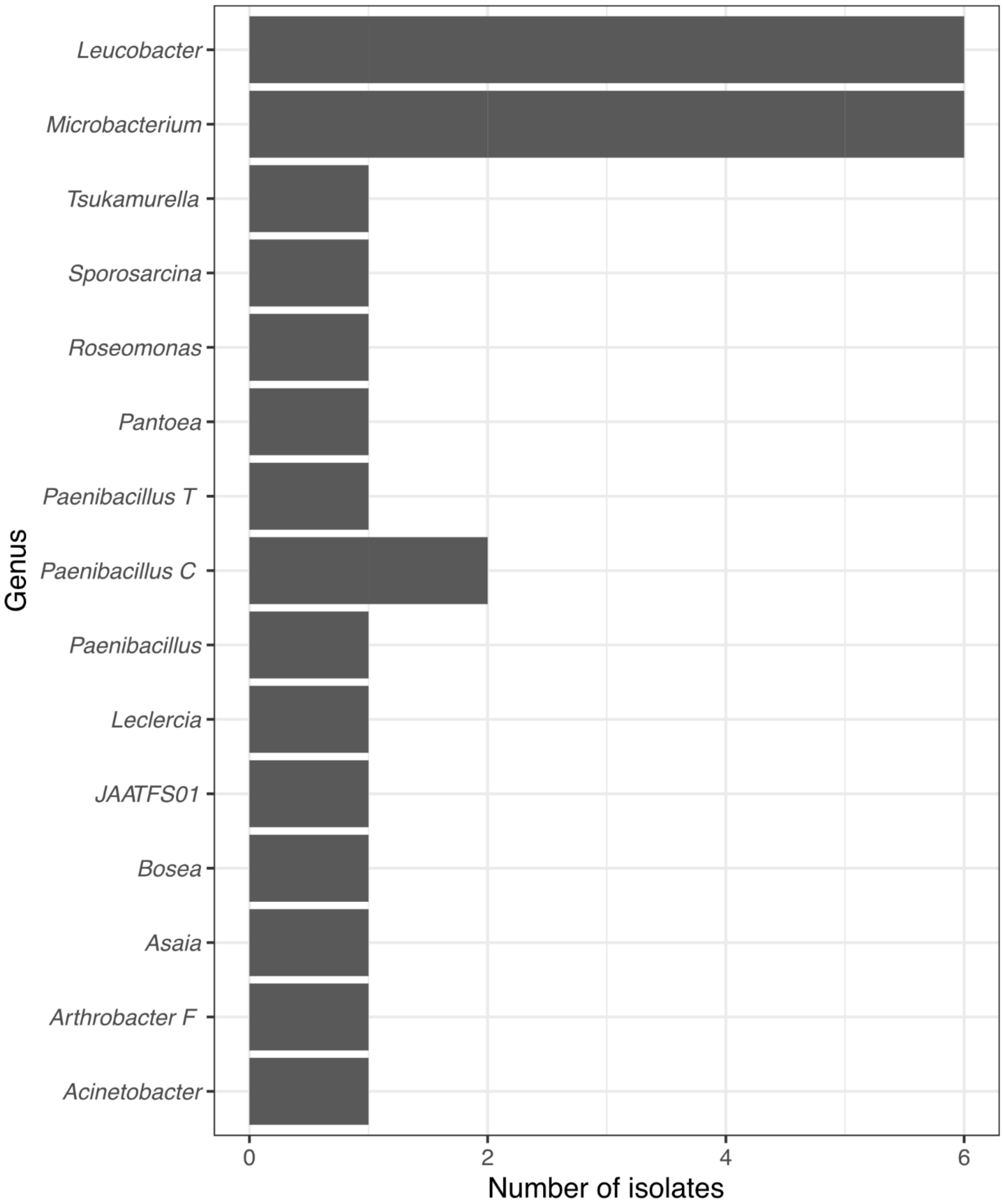


**Fig. S3 | Twenty-five isolates within MosAIC share <95% ANI to a reference genome in the GTDB.** The *x*-axis of the bar chart shows the number of isolates assigned to a reference genome with a given genus assigned taxonomy in the GTDB (*y*-axis). "JAATFS01" is a strain identifier placeholder used by GTDB-Tk when no binomially named representative genome is present in the GTDB.

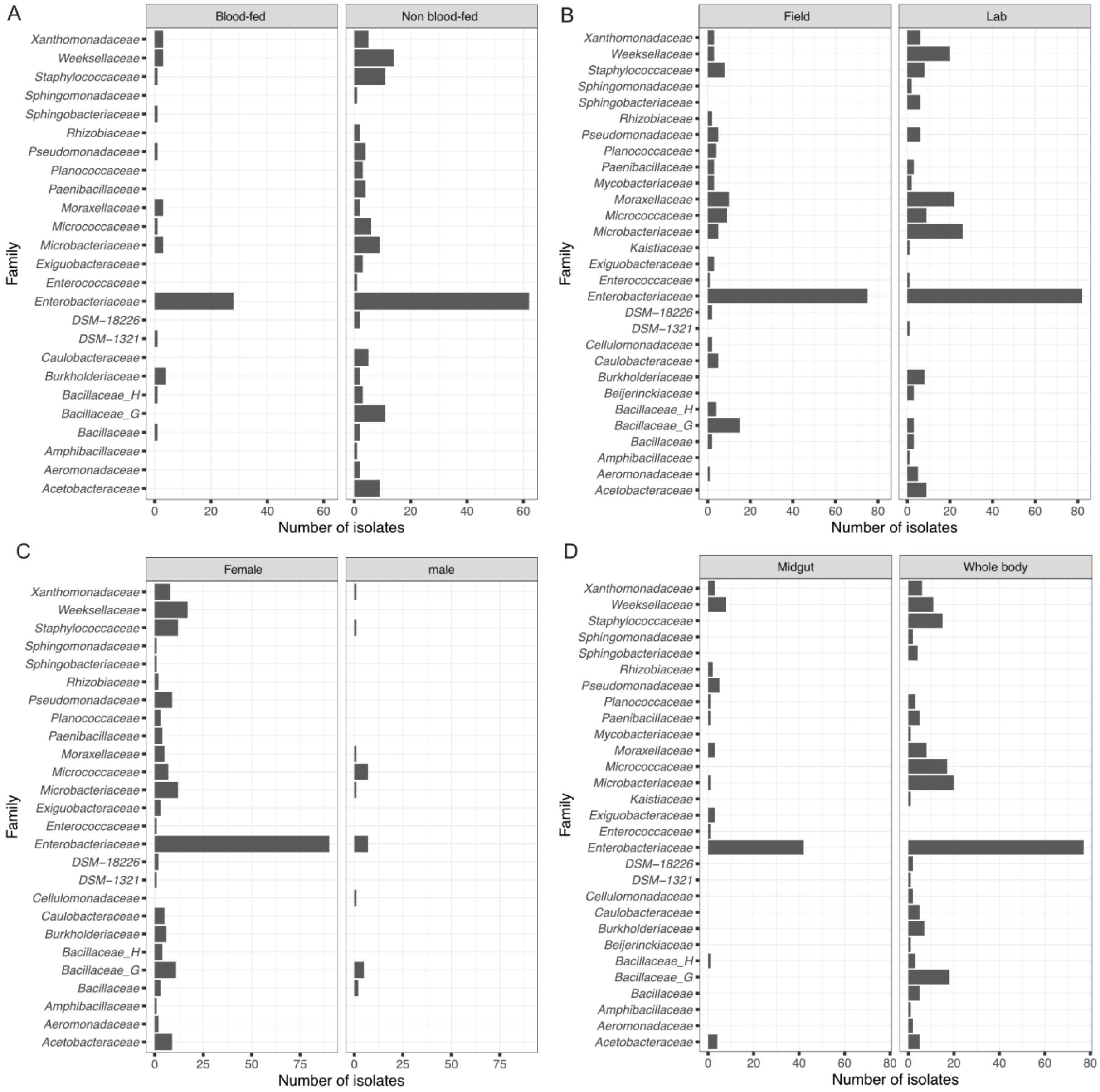


**Fig. S4 | Bacterial family distribution of MosAIC isolates assigned to different metadata categories.** The *x*-axis of each bar chart shows the number of isolates assigned to a reference genome with a given family assigned taxonomy in the GTDB (*y*-axis). Charts are faceted by metadata category as follows: (A) female_feeding_status, (B) lab_field_derived, (C) mosquito_sex, and (D) mosquito_tissue. Metadata category names and definitions follow those presented in Table S1.

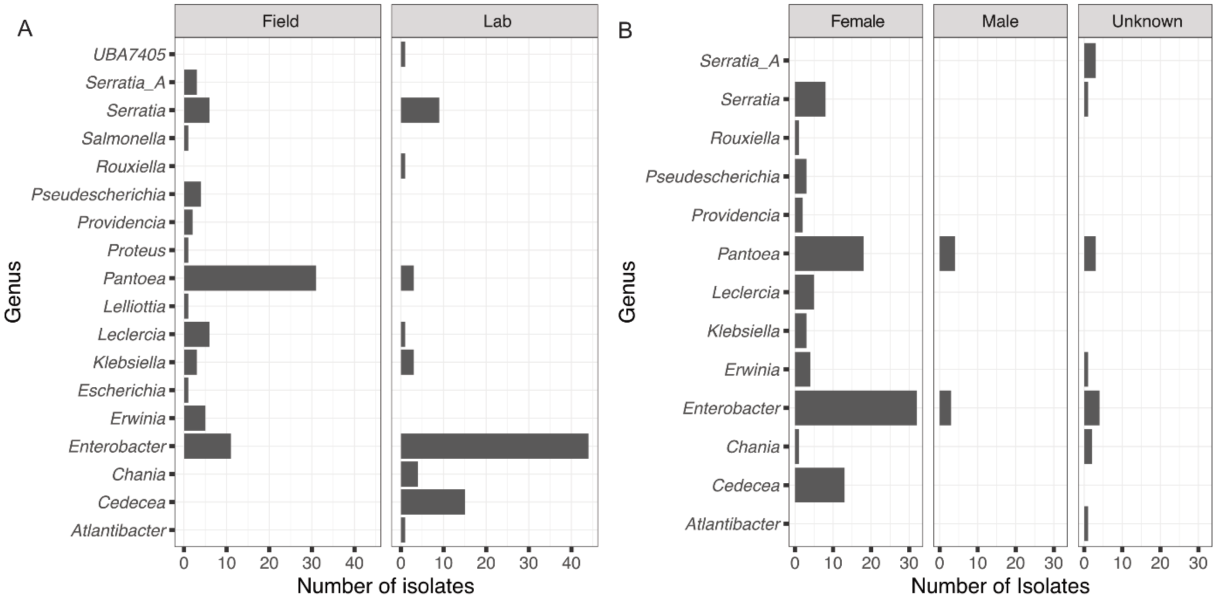


**Fig. S5 | Bacterial genus distribution of MosAIC isolates assigned to different metadata categories.** The *x*-axis of each bar chart shows the number of isolates assigned to a reference genome with a given genus assigned taxonomy in the GTDB (*y*-axis). Charts are faceted by metadata category as follows: (A) lab_field_derived, and (B) mosquito_sex. Metadata category names and definitions follow those presented in Supplementary Table 1. Only isolates assigned to genera within the *Enterobacteriaceae* were included in the analysis.

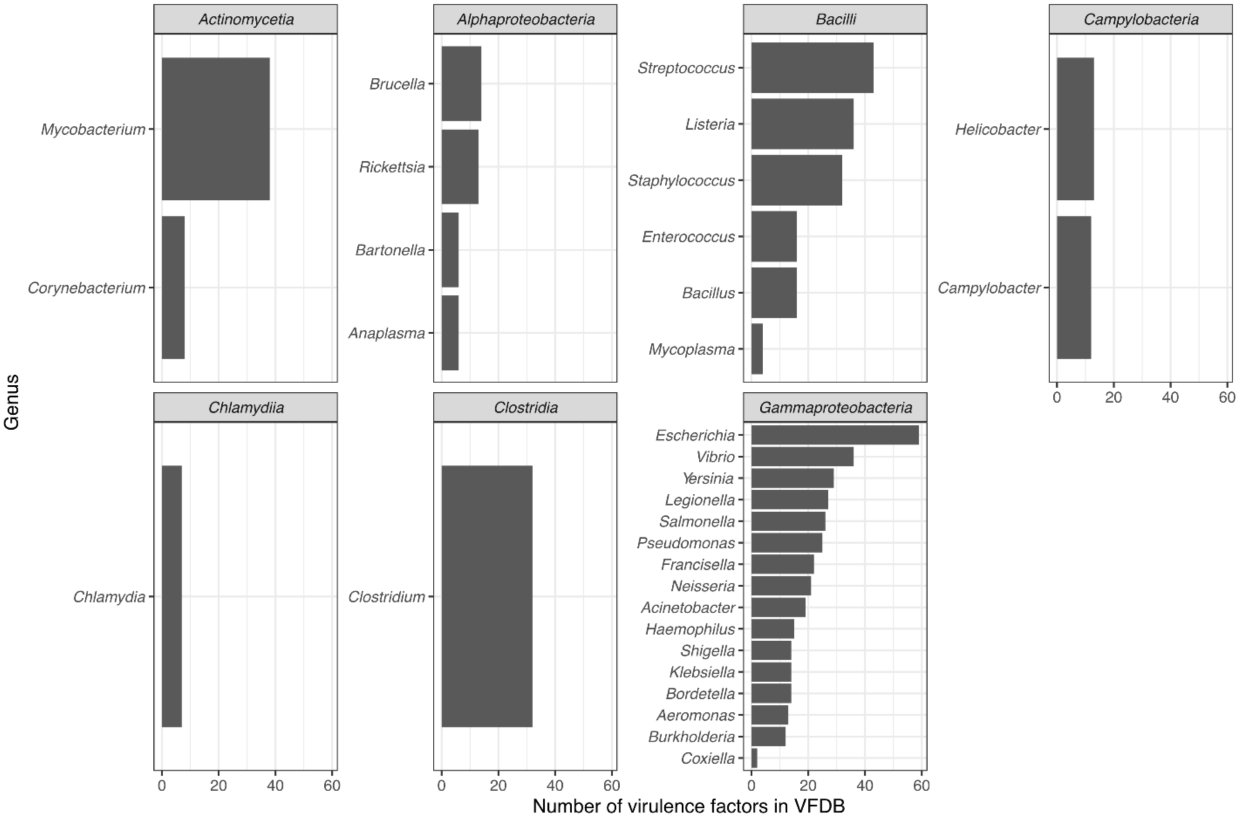


**Fig. S6 | Taxonomic representation across the VFDB.** Charts are faceted by bacterial class, with the *x*-axis of each chart showing the number of isolates assigned to a given genus on the *y*-axis in which at least one virulence factor gene was identified.

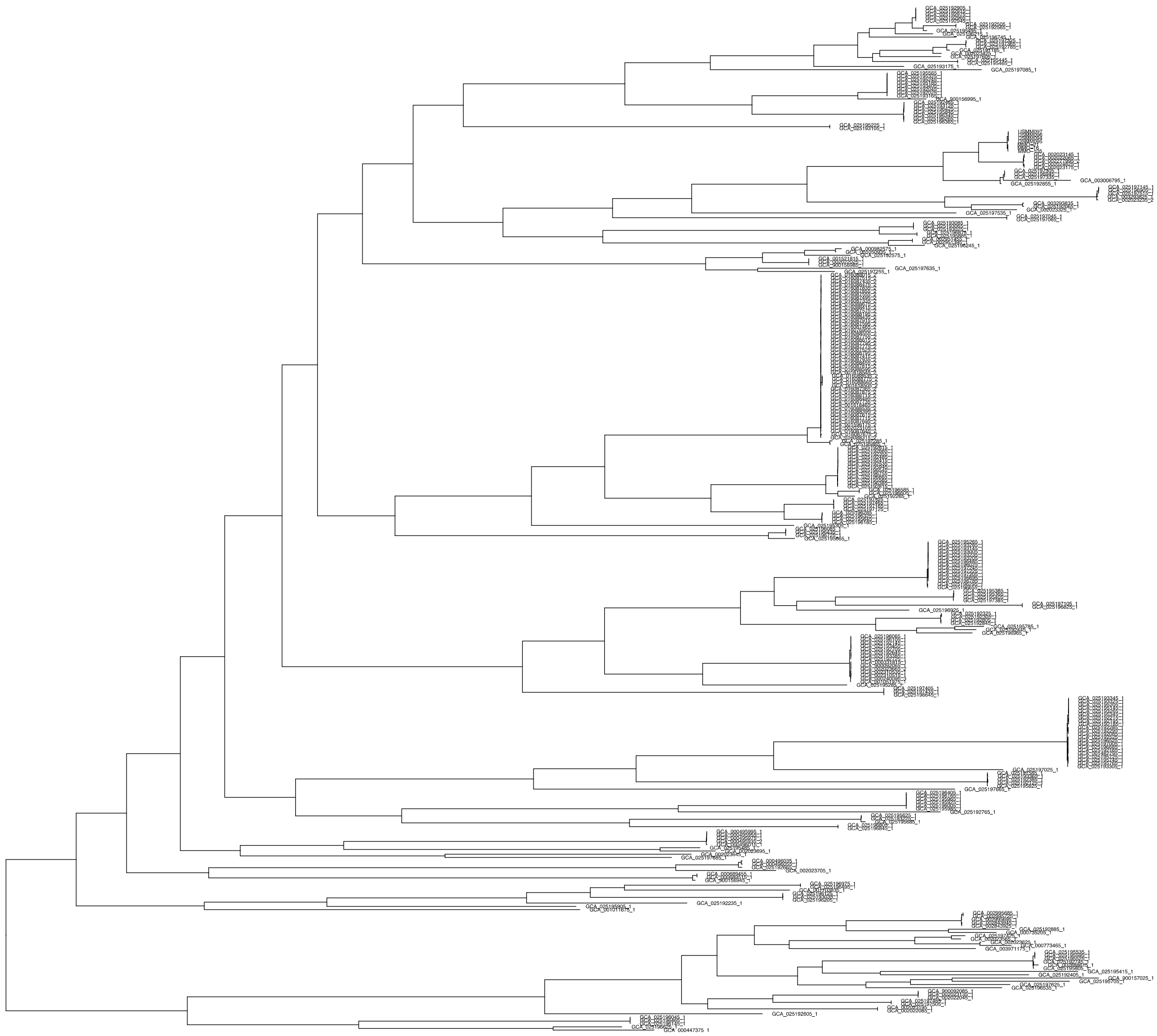

**Fig. S7 | *Elizabethkingia anophelis* Phylogeny with tip labels.** Tip labels of *Elizabethkingia anophelis* shown at each node. Green bar denotes mosquito-associated samples from MosAIC.

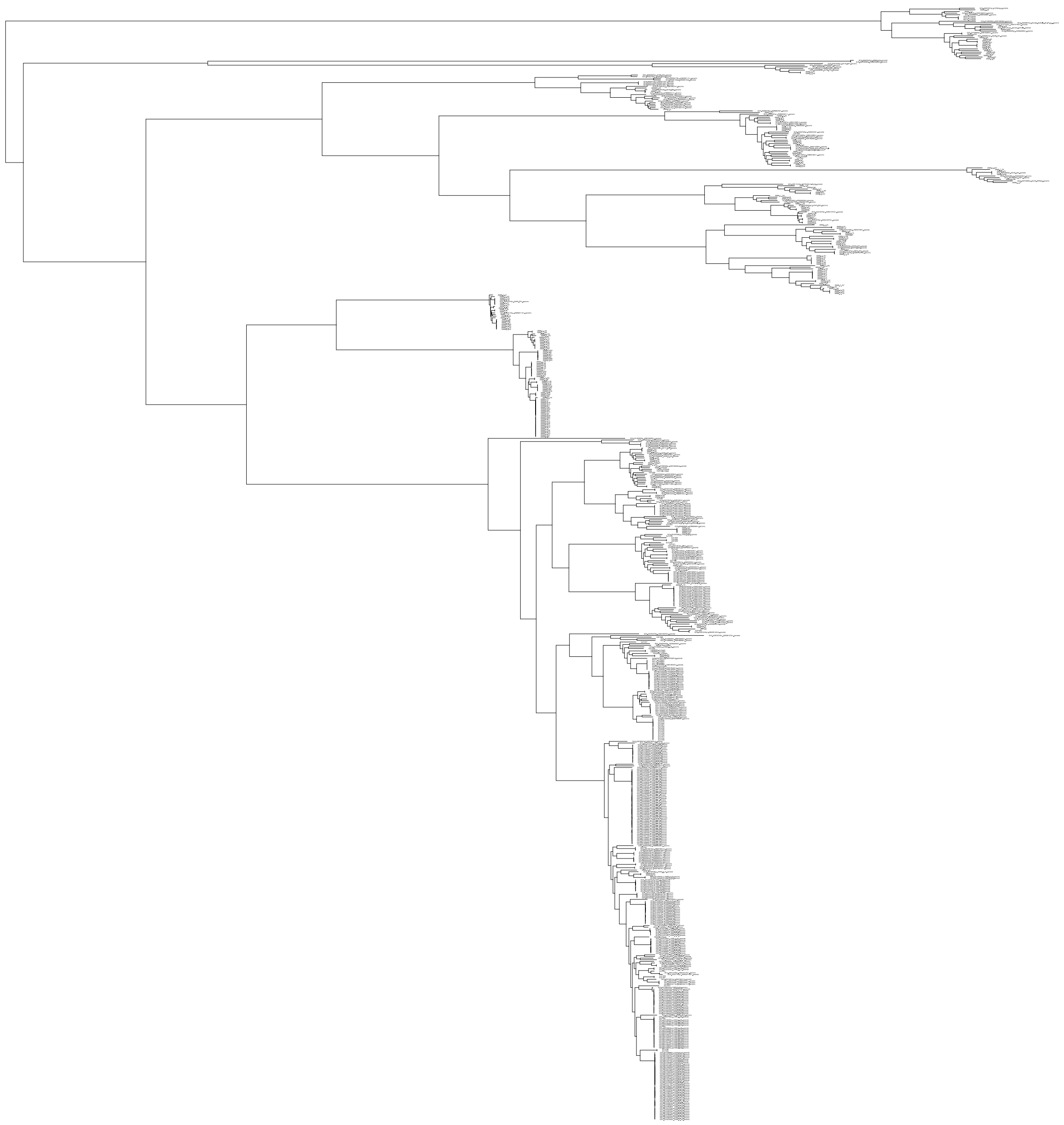

**Fig. S8 | *Serratia* Phylogeny with tip labels.** Tip labels of *Serratia* are shown at each node.

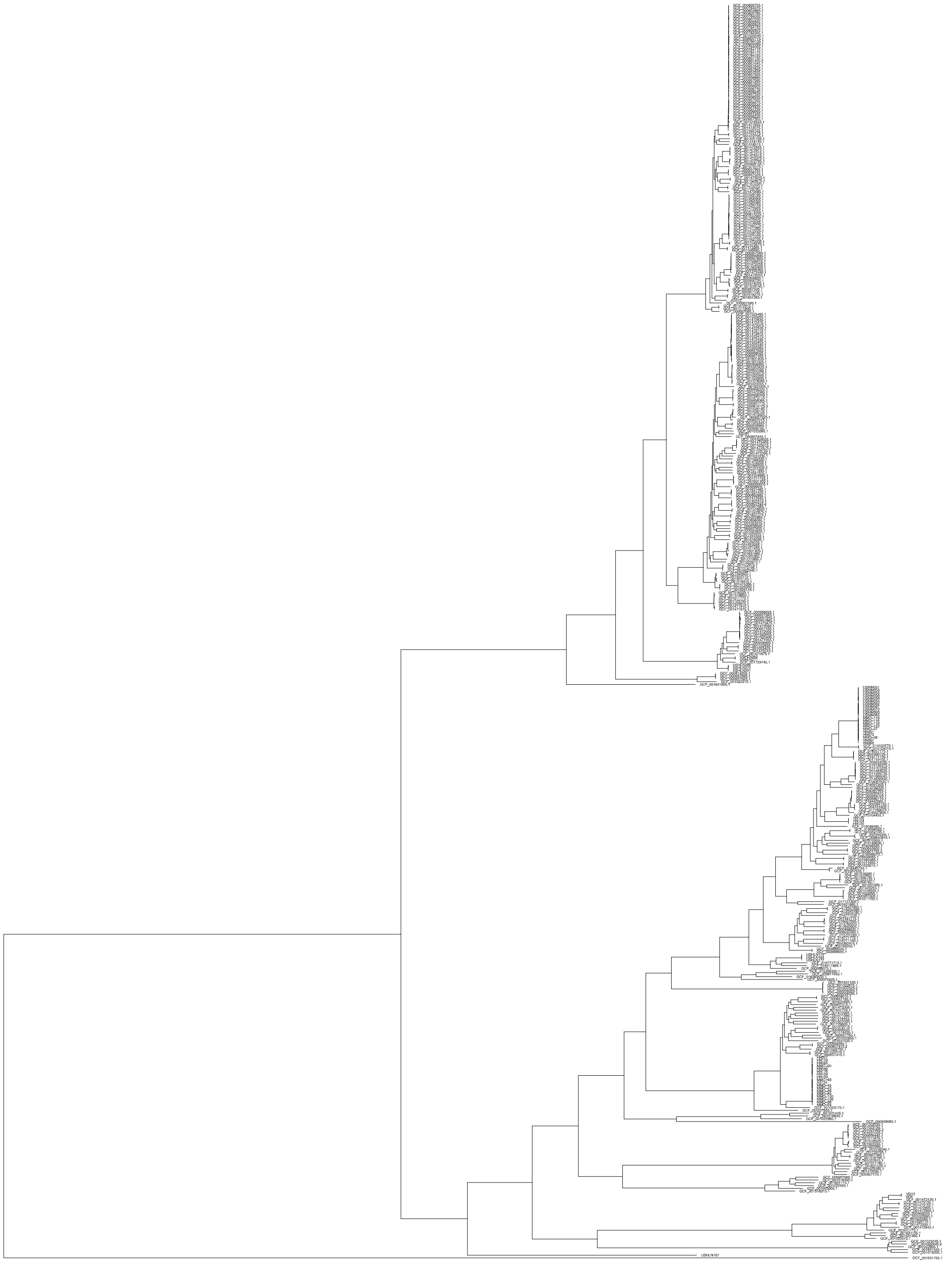

**Fig. S9 | *Enterobacter* Phylogeny with tip labels.** Tip labels of *Enterobacter* are shown at each node.

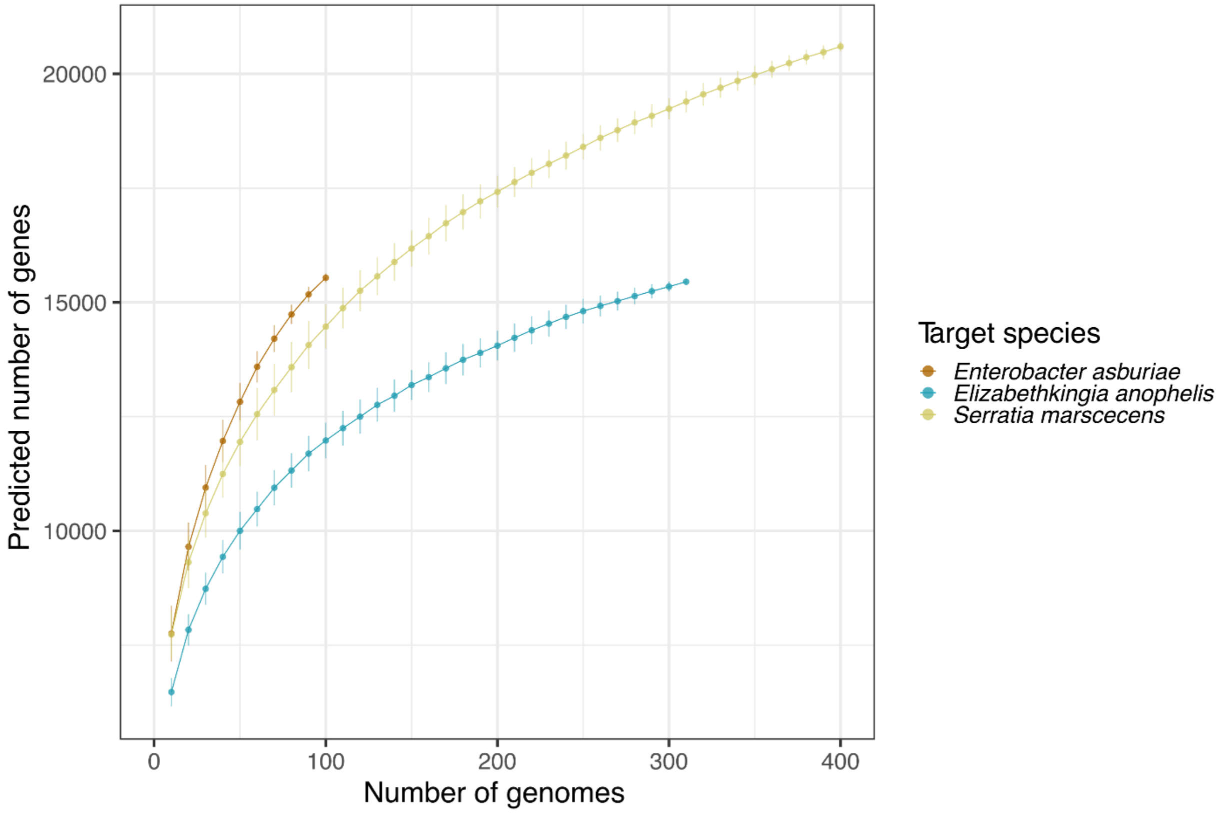


**Fig. S10 | Gene accumulation curve.** Pangenome gene accumulation curve for *En. asburiae, El. anophelis* and *S. marcescens* isolates from MosAIC.

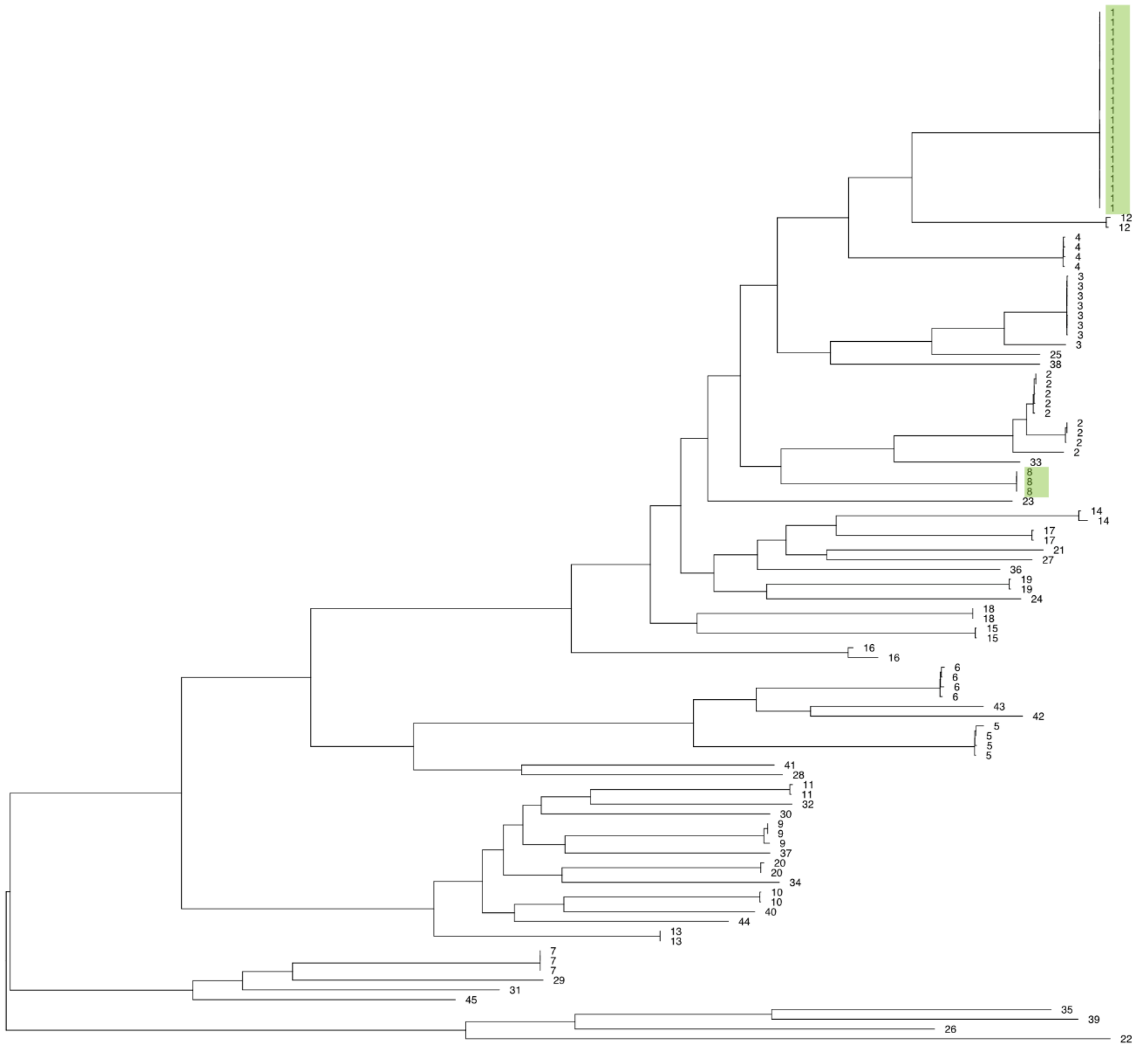

**Fig. S11 | *Enterobacter asburiae* phylogeny overlaid with PopPUNK defined genome clusters.** Tips denote PopPUNK cluster. Green highlight denotes mosquito-associated lineages containing MosAIC isolates.

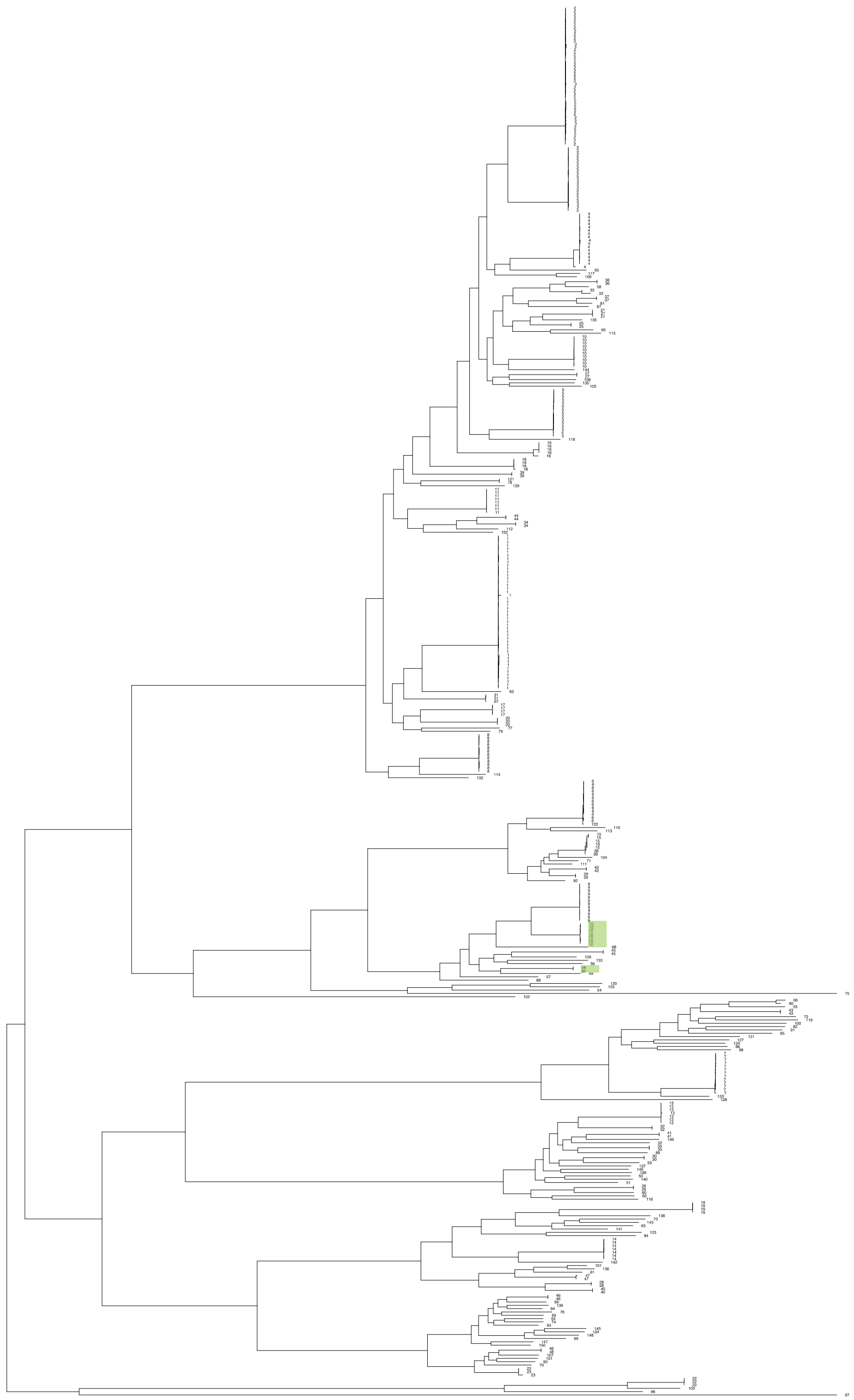

**Fig. S12 | *Serratia marcescens* phylogeny overlaid with PopPUNK defined genome clusters.** Tips denote PopPUNK cluster. Green highlight denotes mosquito-associated lineages containing MosAIC isolates.

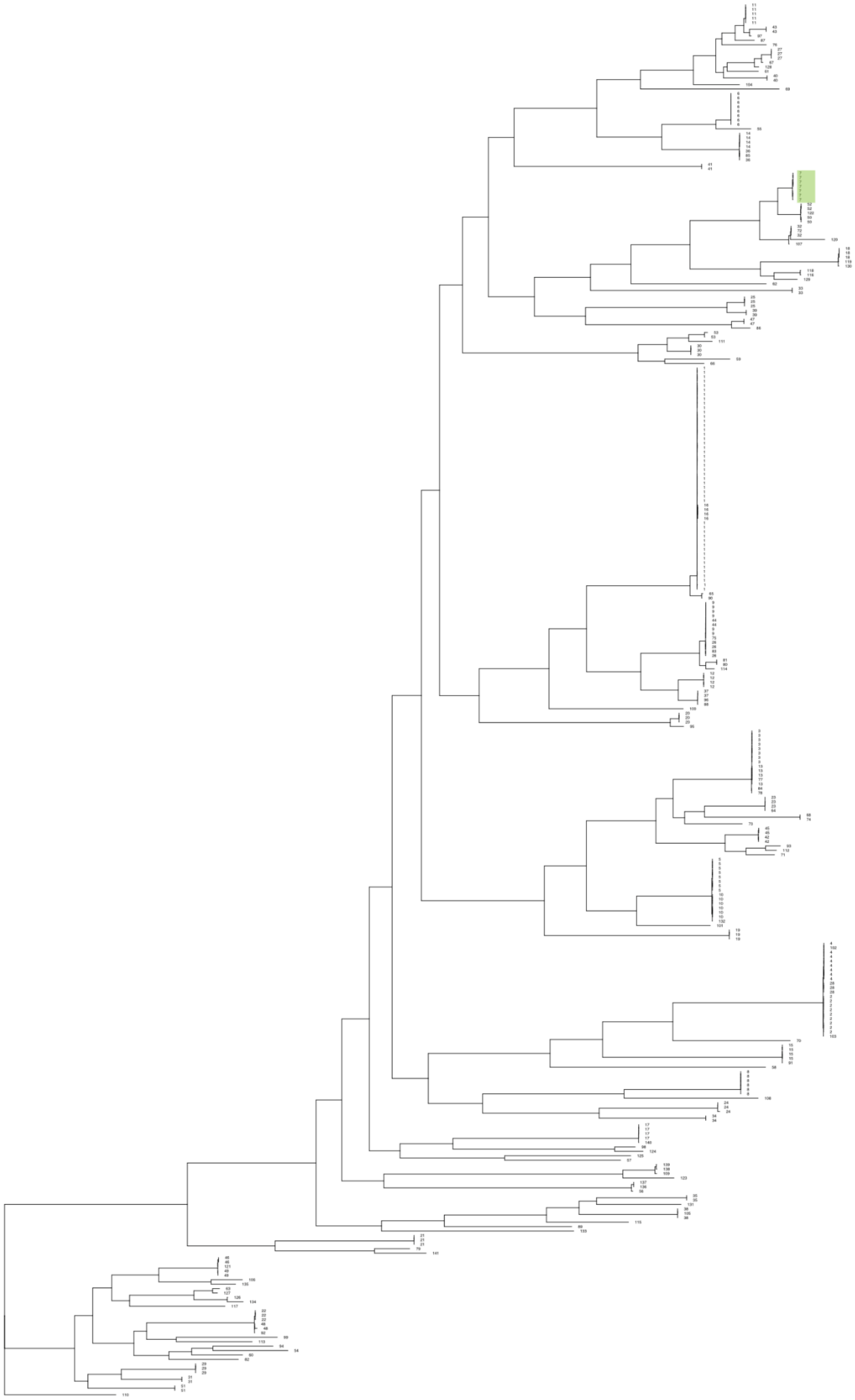

**Fig. S13 | *Elizabethkingia anophelis* phylogeny overlaid with PopPUNK defined genome clusters.** Tips denote PopPUNK cluster. Green highlight denotes mosquito-associated lineages containing MosAIC isolates.

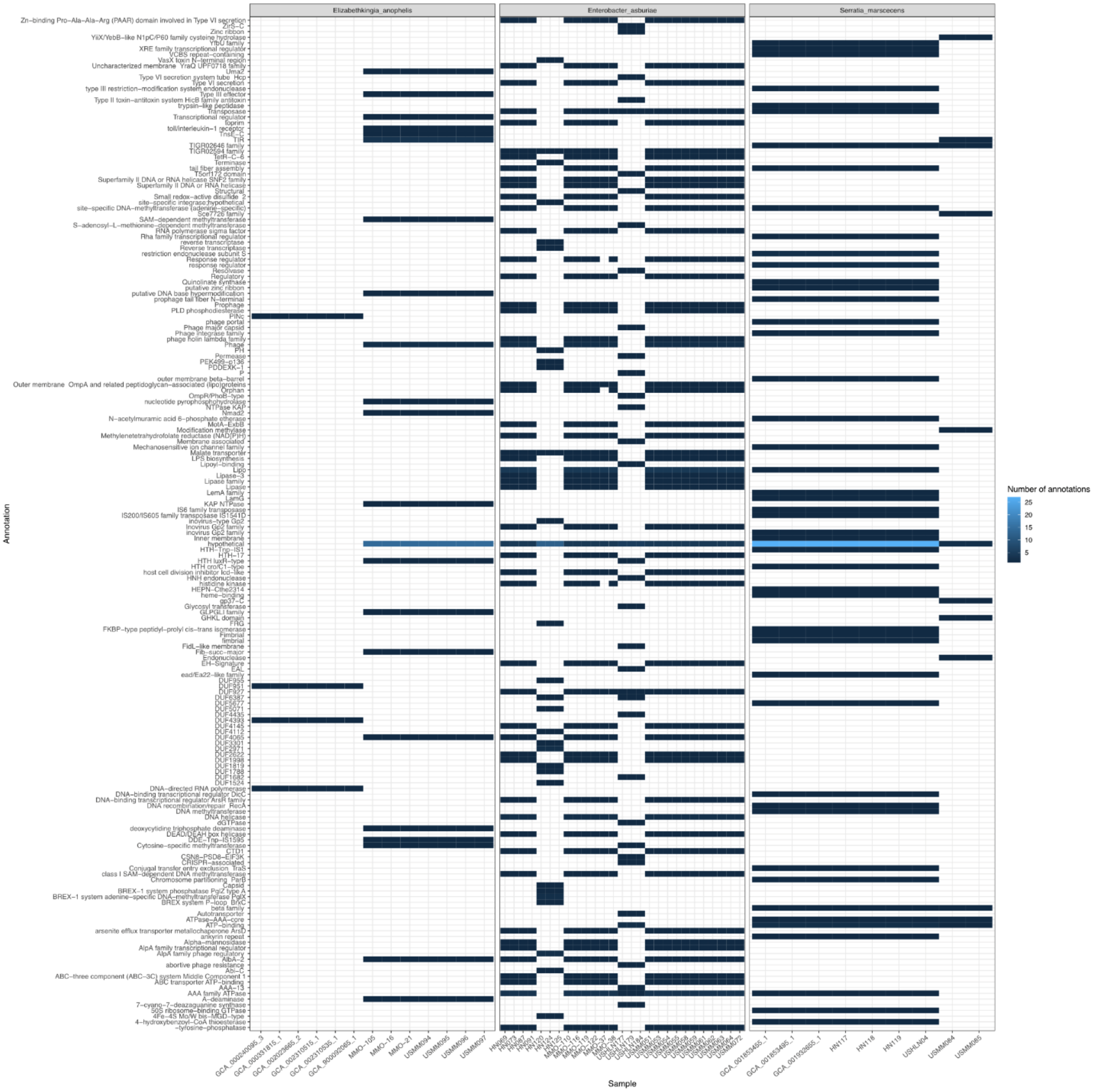


**Fig. S14 | Annotations of lineage-specific core genes identified from mosquito-associated lineages.** Panels summarize annotations for one of three focal species (*Elizabethkingia anophelis, left*; *Serratia marcescens, centre*; or *Enterobacter asburiae, right*), with the *x*-axis of each panel denoting the internal identifier of individual isolates assigned to each species as presented in Table S1. Tiles denote the number of identified annotations corresponding to a given functional category on the *y*-axis, following a gradient from dark blue (few) to light blue (many). White tiles denote categories for which zero annotations were identified in each isolate.

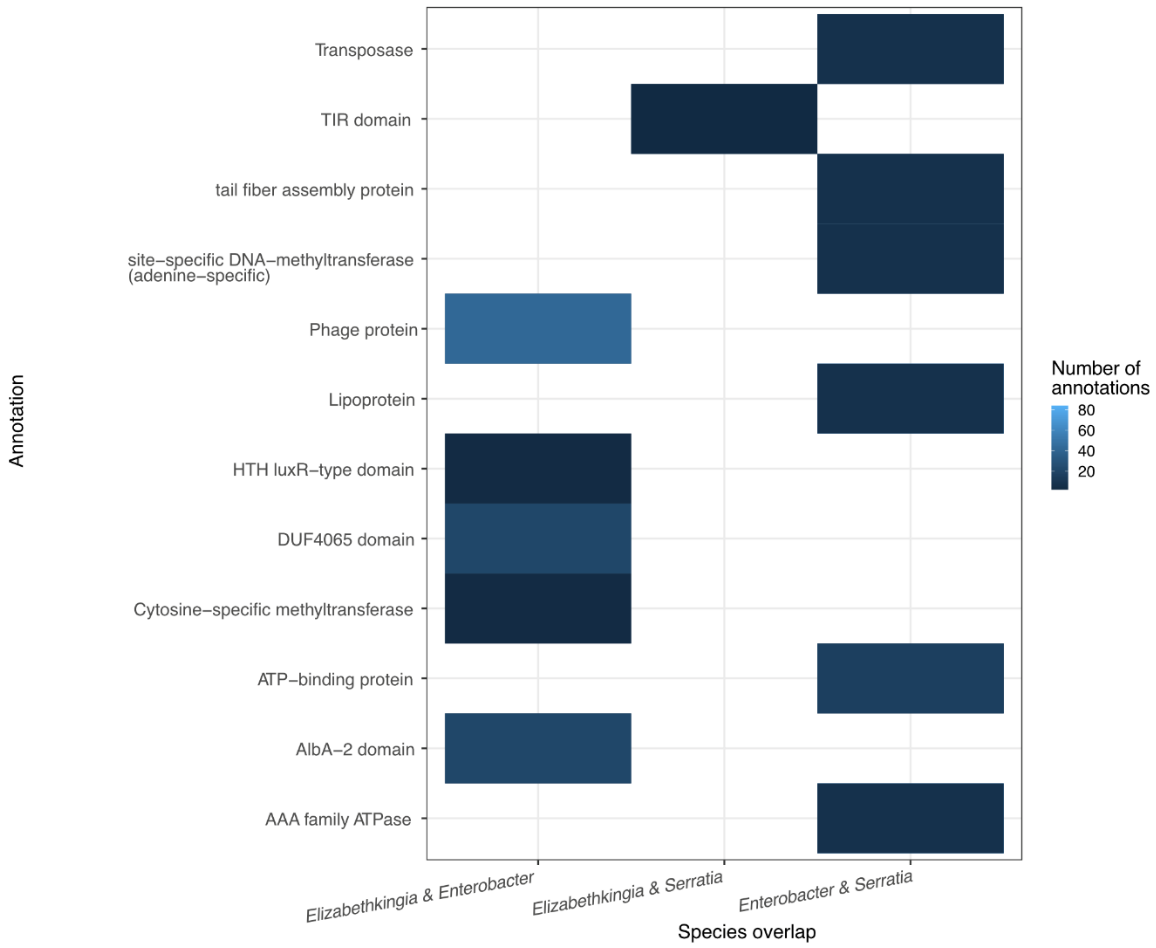


**Fig. S15 | Shared-core gene annotations among mosquito-associated lineages.** Tiles denote the number of identified annotations corresponding to a given functional category on the *y*-axis that were shared between a given species pair on the *x*-axis, following a gradient from dark blue (few) to light blue (many). White tiles denote categories for which zero shared annotations were identified in each species pair.

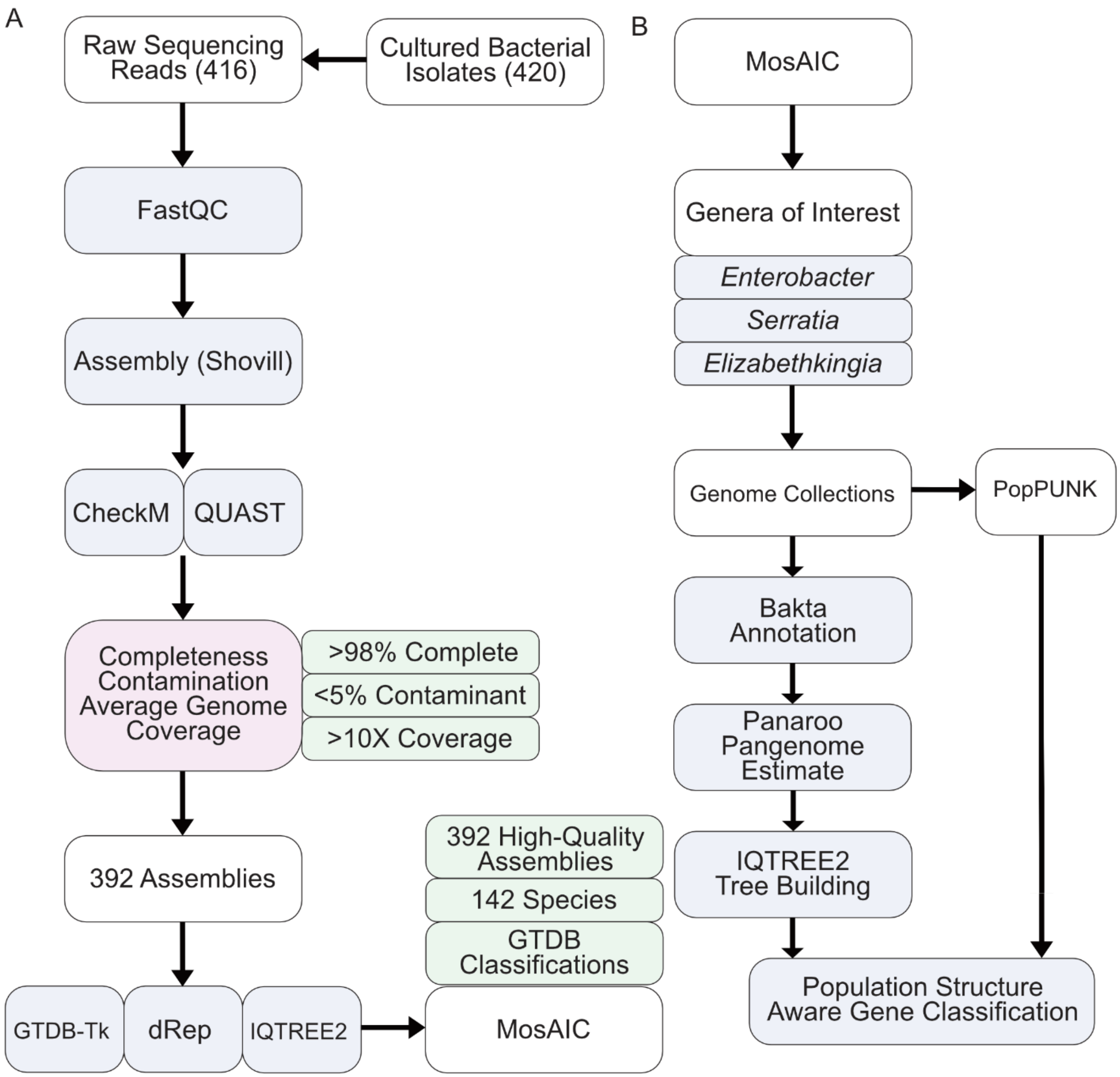


**Fig. S16 | Analysis workflow.** Flowchart describing (A) the assembly of MosAIC genomes and (B) population genomic analyses.
